# Highly divergent and diverse viral community infecting sylvatic mosquitoes from Northeast Brazil

**DOI:** 10.1101/2023.06.27.546706

**Authors:** Alexandre Freitas da Silva, Laís Ceschini Machado, Luisa Maria Inácio da Silva, Filipe Zimmer Dezordi, Gabriel Luz Wallau

## Abstract

Mosquitoes can transmit several pathogenic viruses to humans, but their natural viral community is also composed of a myriad of other viruses such as insect-specific viruses (ISV) and those that infect symbiotic microorganisms. Besides a growing number of studies investigating the mosquito virome the majority are focused on few urban species and relatively little is known about the virome of sylvatic mosquitoes particularly in high biodiverse brazilian biomes. Here, we characterized the RNA virome of 10 sylvatic mosquitos species from Atlantic forest remains at a sylvatic-urban interface in Northeast Brazil employing a metatranscriptomic approach. A total of 16 viral families were detected. The phylogenetic reconstructions of 14 viral families revealed that the majority of the sequences are putative ISVs. The phylogenetic positioning and, in most cases, the association with a high RdRp amino acid divergence from other known viruses suggests that the viruses characterized here represent at least 34 new viral species. Therefore, the sylvatic mosquitoes viral community is majorly composed of highly divergent viruses highlighting the limited knowledge we still have about the natural virome of mosquitoes in general. Moreover, we found that none of the viruses recovered were shared between the species investigated. These findings will help to understand the interactions and coevolution between mosquitoes and viruses in nature.

**Importance:** Mosquitoes are medically important insects as they transmit pathogenic viruses to humans and animals during blood feeding. However, their natural microbiota is also composed of a diverse set of viruses that cause no harm to the insect and other hosts, such as ISVs. In this study, we characterized the RNA virome of sylvatic mosquitoes from Northeast Brazil using unbiased metatranscriptomic sequencing and in depth bioinformatics approaches. Our analysis revealed that these mosquitoes species harbor a diverse set of highly divergent viruses and the majority comprises new viral species. Our findings revealed many new virus lineages characterized for the first time broadening our understanding about the natural interaction between mosquitoes and viruses. Lastly, it also provided several complete genomes that warrants further assessment for mosquito and vertebrate host pathogenicity and their potential interference with pathogenic arboviruses.

## 1. Introduction

Mosquitoes transmit several high impact viral pathogens (arthropod-borne viruses or arboviruses) to humans and animals (Cox, 2010; Manrique-Saide et al., 2010; Wilson and Schlagenhauf, 2016). But besides these pathogenic arboviruses, an increasing number of studies are revealing that mosquitoes hosts a much larger diversity of viruses which are collectively called virome (Atoni et al., 2019a; Pettersson et al., 2019; Shi et al., 2016; Xia et al., 2018). Mosquito viromes are composed by the most abundant and prevalent insect-specific viruses (ISVs) that only infect arthropods (Bolling et al., 2015; de Almeida et al., 2021) and the less prevalent and human pathogenic arboviruses (Vasilakis and Tesh, 2015). Mounting evidence is showing that ISVs may interfere on arboviruses replication either increasing or reducing viral replication in the mosquito body and hence impacting vector competence of specific arboviruses (Bolling et al., 2015; Laureti et al., 2020; Patterson et al., 2020). Some example of ISVs-arbovirus pairs with known interference phenomenon are: Palm Creek Virus - West Nile virus (Hall-Mendelin et al., 2016); Nhumirim virus - Zika virus and Dengue virus 2 (Romo et al., 2018) and Eilat virus - Venezuelan equine encephalitis, Eastern equine encephalitis virus, Chikungunya virus and West equine encephalitis virus (Nasar et al., 2015).

The discovery and characterization of viruses is historically a laborious process requiring cell isolation and classical virology analysis (Mokili et al., 2012; Shi et al., 2018; Zhang et al., 2019). Still, most viruses are not amenable to cell isolation in laboratory conditions limiting our understanding of the natural viral communities to the culturable ones (Shi et al., 2018). But the nucleic acid sequencing revolution of the last decades has opened new possibilities to more comprehensively characterize the natural mosquito viromes. Among several strategies available, bulk metatranscriptome sequencing is one of the less biased approaches for RNA virus genome sequencing (Zhang et al., 2018). Coupled with large scale metagenomic sequencing, new bioinformatic tools are accelerating virus discovery and characterization (Ibañez-Lligoña et al., 2023; Nooij et al., 2018). However, continued development and integration of new tools is required both because of the increasing data volume and the ever increasing difficulty to characterize highly divergent virus genomes (virome dark matter) which compose a large fraction of every virome sequenced (Mokili et al., 2012).

Currently, the majority of mosquito viruses characterized were identified in species from *Culex, Anopheles* and *Mansonia* genera due to a focus on epidemiological important mosquitoes (Atoni et al., 2019a; de Almeida et al., 2021; Moonen et al., 2023). Hence, the large majority of the mosquito species diversity which likely transmit viral pathogens in sylvatic transmission cycles have not been accessed regarding its virome composition. Yet, due to our narrow view of the virosphere and particularly the mosquito viral communities, every new mosquito virome study has revealed many novel viruses. For instance, a recent review of mosquito virome has shown that 14 mosquito genera were positive for viruses which were assigned into 102 viral families (Moonen et al., 2023). In Brazil, there are some scattered studies focusing on characterizing the virome of sylvatic mosquitoes covering different genera such as *Anopheles, Aedes, Culex, Psorophora, Sabethes, Coquillettidia* and *Mansonia* sampled at different the Amazon, Cerrado, Caatinga, Pantanal and Atlantic forest biomes (da Silva et al., 2021a; da Silva Ferreira et al., 2020a; Maia et al., 2019a; Pinto et al., 2017, 2017; Scarpassa et al., 2019a). However, some biomes cover extensive territory and more comprehensive spatiotemporal sampling of mosquitoes are necessary to characterize pathogenic and non-pathogenic components of the natural mosquito viromes.

Here, we performed metatranscriptome sequencing of ten different mosquito species sampled at a sylvatic-urban interface of the Atlantic forest in Northeast Brazil aiming to characterize its RNA virome composition and potential viral threats to humans. We characterized a total of 16 different viral families. The newly discovered viruses exhibit high divergence from known viruses and clustered with previously identified mosquito viruses, but the low amino acid identity at the most conserved gene (RdRp -RNA-dependent RNA polymerase) suggests that they belong to new viral species.

## Material & Methods

### Sample collection and species identification

Mosquito sampling and species identification were performed using the same approach described previously (da Silva et al., 2021b). In brief, we collected mosquitoes using entomological nets during morning and afternoon/evening in two areas of Atlantic forest remains in Northeast Brazil (**Supplementary material 1 - Figure 1**, Parque Estadual Dois Irmãos (PEDI) [8°00′43.3′′S, 34°56′40.7′′W] and Jardim Botânico do Recife (JBR) [8°04′33.0″S, 34°57′35.9″W], Recife municipality, state of Pernambuco, Brazil). The collected mosquitoes were transported alive to the Entomology Department of the Aggeu Magalhães Institute (IAM-FIOCRUZ) and stored at −80°C until the taxonomic identification. Collected mosquito samples were taxonomically identified using dichotomous keys available in the literature for neotropical Culicidae (Forattini, 2002). All specimens were processed in a cooled bench (∼0°C) to restrict RNA degradation. After morphological identification, each specimen had its abdomen dissected and processed for DNA extraction using the protocol from Ayres et al. (2003), while the remaining tissues were stored in the −80°C until species-specific pooling and the RNA extraction procedures. We amplified the Cytochrome c oxidase subunit I (COI) for individual abdomen DNA using the GoTaq (Promega) protocol, and carried out sequencing using the Sanger method on an ABI 3500xL (Applied Biosystems), following the manufacturer’s instruction. Electropherograms were analyzed on Geneious Prime® 2020.0.5 to extract the consensus fasta sequence. A BLASTn analysis (Altschul et al., 1990) was carried out with these sequences against a custom database composed by NCBI and BOLD mosquito COI sequences (retrieved on March 4th 2020), and also a database of mitochondrial genomes from local mosquitoes from da Silva et al., (2020) to confirm and ensure the mosquito species used in the pooling scheme were from the same taxon. The only exception was the *Aedes albopictus* samples that were only identified by morphological characters due the ease visualization of species-specific morphological characters. The reconstructed phylogenetic tree of the COI sequences corroborated previous database similarity searches (**Supplementary material 1 - Figure 2**).

### RNA extraction and sequencing

Based on previous mosquitoes identification, the remaining tissues stored at −80°C from the same taxon were pooled in ultrapure water and macerated using a pestle motor tissue grinder. One hundred microliter (µl) of macerated samples were used to extract the total RNA following the Trizol protocol as suggested by the fabricant (Invitrogen, USA). The pellet was eluted in a final volume of 30 ul. The extracted nucleic acids were treated with TURBO™ DNase (2u/µl - Ambion) following the manufacturer’s instructions. RNA samples were quantified and quality checked through Qubit RNA HS kit and Bioanalyzer respectively. Total RNA samples were processed for ribosomal RNA depletion with the RiboMinus™ Eukaryote System v2 kit following manufacturer’s instructions (Invitrogen, USA). The sequencing library was prepared using the TruSeq Stranded Total RNA library kit (Illumina, USA) and sequenced on a NextSeq 500 Illumina platform using a paired-end approach of 75 bp.

### Virome characterization and taxonomic classification

The viromes were characterized following the study workflow presented in **Supplementary material 1 - Figure 3**. Sequenced reads were firstly quality checked using FastQC tool (https://www.bioinformatics.babraham.ac.uk/projects/fastqc/) followed by trimming of low quality reads using Trimmomatic (Bolger et al., 2014) with the following parameters: LEADING:3 TRAILING:3 SLIDINGWINDOW:4:20 MINLEN:36 and TruSeq2-PE as the adapters file to be removed. The HTML reports from FastQC tool were parsed using the MultiQC tool (Ewels et al., 2016) that combined the data in a single report used for general genomic statistics recovery. The metatranscriptome assemblies were generated using three different assembly tools: Trinity v.2.11 (Grabherr et al., 2011), rnaSpades (Bushmanova et al., 2019) and metaSpades (Nurk et al., 2017) in default mode. This approach was used to obtain a more complete assembly as shown by previous studies previously (Roux et al., 2017; Sutton et al., 2019). All contigs obtained from the three different assembly tools described above were clustered using the CD-HIT-EST tool (Li and Godzik, 2006) with the following parameters: -c 0.98 -G 0 -aS 0.9 -g 1 -n 9, in order to exclude redundant sequences having >= 98% of identity and retain the larger contigs. In order to identify and recover viral sequences from the assembled metatranscriptomes, we firstly download protein sequences assigned with the viral taxonomy tag (txid10239) from the National Center for Biotechnology Information (NCBI) on December 21th, 2020. The sequences were clustered using the CD-HIT tool (Li and Godzik, 2006) with the following parameters: -G 0 -aS 0.95 -g 1 -M 100000 -n 5 aiming to exclude the redundant sequences. Based on the assembled metatranscriptome we predicted the amino acid sequences encoded from contigs using the Prodigal tool (Hyatt et al., 2010) employing the meta flag. Then, we identified the viral contigs using DIAMOND (Buchfink et al., 2015) performing a blastp search of the predicted amino acid sequences against the retrieved viral protein database (txid10239) and another four additional searches were performed using specific databases of viral RdRp sequences such as NeoRdRp (Sakaguchi et al., 2022), PalmDB (Edgar et al., 2022), RVMT (Neri et al., 2022) and RdRp-Scan (Charon et al., 2022). These databases were used aiming to recover divergent and unclassified viruses that are not presented in the NCBI txid10239 database. False positive viral hits were removed through a second DIAMOND blastp search against the non-redundant protein database (NR) from NCBI (retrieved on December 12th, 2023). Only contigs where the first 5 best hits of the predicted amino acid sequences that showed similarities with viral proteins in both DIAMOND analysis were kept for further analysis. All DIAMOND analyses were run using the E-Value of 1E-3 and using the more-sensitive parameter.

To further in depth characterize more distant related viral contigs that might have been missed from the DIAMOND search, we performed another approach of viral identification based on Hidden Markov models (HMM) profiles using HMMER3 (http://hmmer.org/). The predicted amino acid sequences were used in the hmmsearch analysis comparing it with HMM models available from the RVDB-prot-HMM database and those HMM models available for the previously cited databases (RVMT, RdRp-Scan and NeoRdRp). Hits showing E-Value of 1E^-3^ were selected for further analysis.

In order to exclude endogenous viral elements (EVEs) that are actively transcribed from the mosquito genomic loci we performed a BLASTn search using the previously identified viral contigs (DIAMOND and HMM analysis) as queries against mosquito genomes database (**Supplementary material 1 -Table 2** and **Supplementary material 1 - Table 3**). Firstly we download all available mosquito genomes from NCBI (retrieved on September 9th 2023, **Supplementary material 1 - Table 2**) and we combined with draft whole mosquito genomes sequenced on Hiseq or Novaseq Illumina technology for the studied species (**Supplementary material 1 - Table 3**). These draft mosquito genomes used here were assembled using the megahit tool (Li et al., 2015). The viral contigs showing hits with genomic regions containing a minimum query coverage of 50% and more than 90% of identity were excluded from further analysis as similarly performed by other authors (Palatini et al., 2022).

To obtain the virome profile for the studied mosquito species, we filtered the contigs larger than 600 bp and we used the taxonomic annotation from the set of sequences containing RdRp hallmark protein, the most conserved and used marker to viral classification. The best hit from DIAMOND and HMM analysis for each query were used to obtain the viral contigs taxonomy information. Since RdRp naming in both NCBI sequences or RVDB-hmm-prot profiles can vary, we performed searches for different terms aiming to filter the RdRp sequences from previous analysis (**Supplementary material 1 - table 1**). Taxonomic information recovered from DIAMOND was used when sequences showed taxonomic information from both methodologies (DIAMOND and HMM analysis). Sequences showing best hits with unclassified viruses were phylogenetically analyzed and reclassified based on annotation of the closest homologs as long as it clustered in monophyletic clades with high node support.

In order to obtain the number of reads per viral contig we mapped quality-trimmed reads against the identified viral contigs using Bowtie2 in default mode and statistical metrics were collected using coverM tool (https://github.com/wwood/CoverM). The raw reads were submitted to the European Bioinformatic Institute under the project number: PRJEB63303 and Run Accessions: ERR12411109-ERR12411118.

In order to evaluate viral genome/segment completeness of the identified viral contigs we used the ViralComplete module from Metaviralspades tool (Antipov et al., 2020). Briefly, the ViralComplete analysis predicts the genes and the viral proteins from the contigs and performs a BLAST analysis against the viral RefSeq databases. Moreover, the tool compares if the length of analyzed contig is similar to the viral hit and then classifies them into full-length or partial sequences.

### Phylogenetic analysis

Phylogenetic trees were reconstructed for each viral family that showed contigs encoding RdRp sequences larger than 600 bp and amino acid sequences larger than 100 aa. We included in the analysis approximately fifty best hit sequences recovered from DIAMOND blastp against the NR database. The viral proteins were aligned using MAFFT v.7 (Katoh and Standley, 2013) with default parameters. The non-aligned blocks were removed from the alignment using the Trimal tool (Capella-Gutiérrez et al., 2009). The phylogenetic trees were reconstructed using the IQ-TREE 2.0 (Nguyen et al., 2015) and the best evolutionary model for each alignment was selected by the ModelFinder (Kalyaanamoorthy et al., 2017) implemented in the same tool. The node supports were evaluated with 1,000 replicates of ultrafast bootstrapping (Hoang et al., 2018). The consensus trees were visualized and annotated with the *ggtree* (Yu et al., 2017) package from R programming language. The genomic maps displayed on trees were built using the *gggenomes* R package (https://github.com/thackl/gggenomes).

The phylogenies of mitochondrial genomes and COI from mosquitoes were reconstructed using the same approach described above. In brief, draft mitochondrial genomes for each mosquito species were assembled using the metatranscriptome data on MITObim 1.9 (Hahn et al., 2013). Then, nucleotide sequences were aligned using MAFFT v.7 and trimmed using the Trimal tool. The phylogenetic tree was obtained and plotted using IQ-TREE 2.0 and Figtree v1.4.4 (http://tree.bio.ed.ac.uk/software/figtree/) respectively.

## Results

A total of 194 specimens were processed and identified at species level. The mosquito species were splitted into ten pools representing the following species: *Mansonia titillans, Mansonia wilsoni, Ae. albopictus, Psorophora ferox, Aedes scapularis, Coquillettidia chrysonotum, Coquillettidia. venezuelensis, Limatus durhamii, Coquillettidia hermanoi, Coquillettidia albicosta* (**Supplementary material 1 - Figure 1** and **Supplementary material 1 - Table 4**).

The Illumina sequencing generated a total of 88.99 Gbp representing 593.3 million of paired end reads distributed along the ten mosquito pools (**Table 1)**. The generated reads per pools ranged from 47.1 million of paired reads for *Ae. scapularis* to 69.1 million of paired reads for *Cq. chrysonotum* (**Table 1)**. A total of 1,971,030 contigs were generated from the three assembly approaches (Trinity, metaSpades and rnaSpades), while 1,285,976 were retained after redundancy removal (**Table 1)**.

**Table 1.** General sequencing and viral identification statistics.

| Species name | Library name | Million of Raw paired reads | Million of trimmed paired reads* | Total contigs** | Total clustered contigs | Total viral contigs*** | Total viral reads | Full-length viral seqs**** |
| --- | --- | --- | --- | --- | --- | --- | --- | --- |
| *Cq. venezuelensis* | P2 | 65.40 | 61.8 | 161,589 | 99,069 | 14 | 7,311 | 4 |
| *Ma. wilsoni* | P8 | 50.20 | 47.1 | 204,403 | 129,918 | 13 | 89,143 | 5 |
| *Cq. albicosta* | P1 | 61.30 | 57.1 | 263,504 | 178,387 | 10 | 21,406 | 1 |
| *Ma. titillans* | P10 | 68.40 | 64.4 | 210,612 | 132,164 | 9 | 748,526 | 3 |
| *Li. durhamii* | P5 | 51.40 | 48.5 | 139,978 | 80,031 | 7 | 48,365 | 0 |
| *Cq. chrysonotum* | P6 | 69.50 | 65.6 | 175,237 | 115,351 | 3 | 18,428 | 1 |
| *Cq. hermanoi* | P9 | 58.80 | 54.7 | 289,556 | 192,588 | 3 | 207 | 0 |
| *Ae. albopictus* | P7 | 69.10 | 64.5 | 260,363 | 178,599 | 2 | 49,007 | 2 |
| *Ae. scapularis* | P3 | 47.10 | 44.3 | 139,261 | 90,437 | 2 | 13,633 | 2 |
| *Ps. ferox* | P4 | 52.10 | 48.5 | 126,527 | 89,432 | 1 | 3,173 | 0 |
| Total | - | 593.30 | 556.5 | 1,971,030 | 1,285,976 | 64 | 999,304 | 18 |

\*Number considering the paired reads plus the unpaired. \*\* Considering Trinity, rnaSpades and metaSpades assemblies. \*\*\* Considering only those viral contigs with RdRp markers. \*\*\*\* Analysis from ViralComplete.

After performing DIAMOND searches using the viral protein database (NCBI taxid10239) we were able to identify sixty one viral contigs (**Figure 1A**). The additional searches based on RdRp databases were able to identify a total of fifteen viral contigs (**Supplementary material 1 - Material 1 - Figure 4A and B**). The majority of viral contigs were identified by all four RdRp databases (**Supplementary material 1 - Figure 4B**). However in some cases only two (RDRP-SCAN and RVMT) or three (NEORDRP, RVMT and RDRP-SCAN) databases have identified specific viral contigs (**Supplementary material 1 - Figure 4B**). NEORDRP database was the only one that could identify viral contigs not identified by the other databases (**Supplementary material 1 - Figure 4B**). Comparing the viral contigs identified using RdRp databases together with those identified from NCBI taxid10239 database, we obtained a total of 63 viral contigs (**Figure 1A**). The majority of them were identified uniquely by the NCBI taxid10239 database and a minority using the RdRp databases (**Figure 1A**). Two viral contigs were only identified by RdRp databases using NEORDRP, RVMT and RDRP-SCAN (**Figure 1A**). The HMM analysis was able to identify a total of sixteen viral contigs and the majority were identified by the all four HMM model datasets (**Figure 1B**). No viral contig was uniquely identified by one specific HMM model dataset. Comparing the viral contigs identified by both approaches, among the sixteen identified using HMM analysis, fifteen were also identified using DIAMOND analysis and only one was uniquely identified using HMM models (**Supplementary material 1 - Figure 4C**).

**Figure 1.**
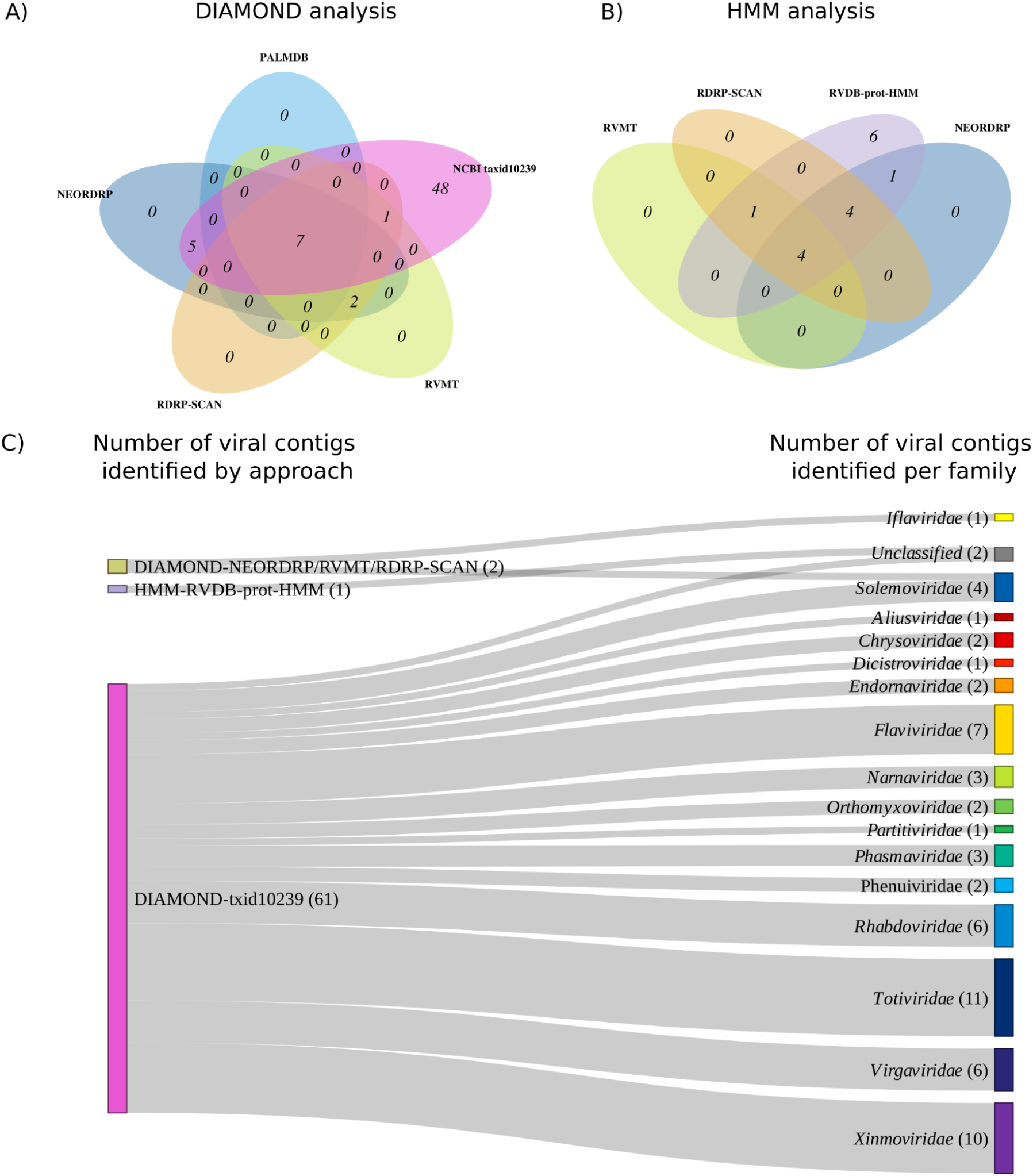
General information on viral identification. A) Venn diagram illustrating the overlap of viral contigs identified using different viral databases on DIAMOND pairwise alignment approach. B) Venn diagram illustrating the overlap of viral contigs identified using different HMM models search. C) Sankey diagram showing the viral families identified by the different approaches employed on viral identification.

A total set of 64 viral contigs were identified using the two approaches (DIAMOND and HMM analysis) and we were able to identify sixteen viral families (**Figure 1C** and **Figure 2D-E**). The majority of viral families (*Aliusviridae, Chrysoviridae, Dicistroviridae, Endornaviridae, Flaviviridae, Iflaviridae, Narnaviridae, Orthomyxoviridae, Partitiviridae, Phasmaviridae, Phenuiviridae, Rhabdoviridae, Totiviridae, Virgaviridae, Xinmoviridae*) were identified using only DIAMOND and NCBI taxid10239 database, while *Solemoviridae* was identified by NCBI taxid10239, NEORDRP, RVMT and RDRP-SCAN databases (**Figure 1C**) and *Ifaviridae* was identified only by NEORDRP, RVMT and RDRP-SCAN. Only one viral contig was identified using HMM analysis and it was not able to be classified (**Figure 1C**). The majority of the viral contigs were classified into known viral families (**Figure 2A**). Three viral genomic architectures were found analyzing all dataset of identified viral contigs: the majority belonged to ssRNA(-), followed by ssRNA(+) and few viral contigs were annotated as dsRNA (**Figure 2C** and **Supplementary file 1**). Furthermore, we recovered a total of eighteen full-length viral sequences from segmented viruses or linear complete genome from eight viral families (*Orthomyxoviridae, Partitiviridae, Phenuiviridae, Rhabdoviridae, Solemoviridae, Totiviridae, Virgaviridae* and *Xinmoviridae*) (**Figure 2B** and **Table 1**), while the remaining sequences were classified as partial. The full-length sequences ranged from 1.75 kb to 13.08kb (**Supplementary material 1 - Figure 5**).

**Figure 2.**
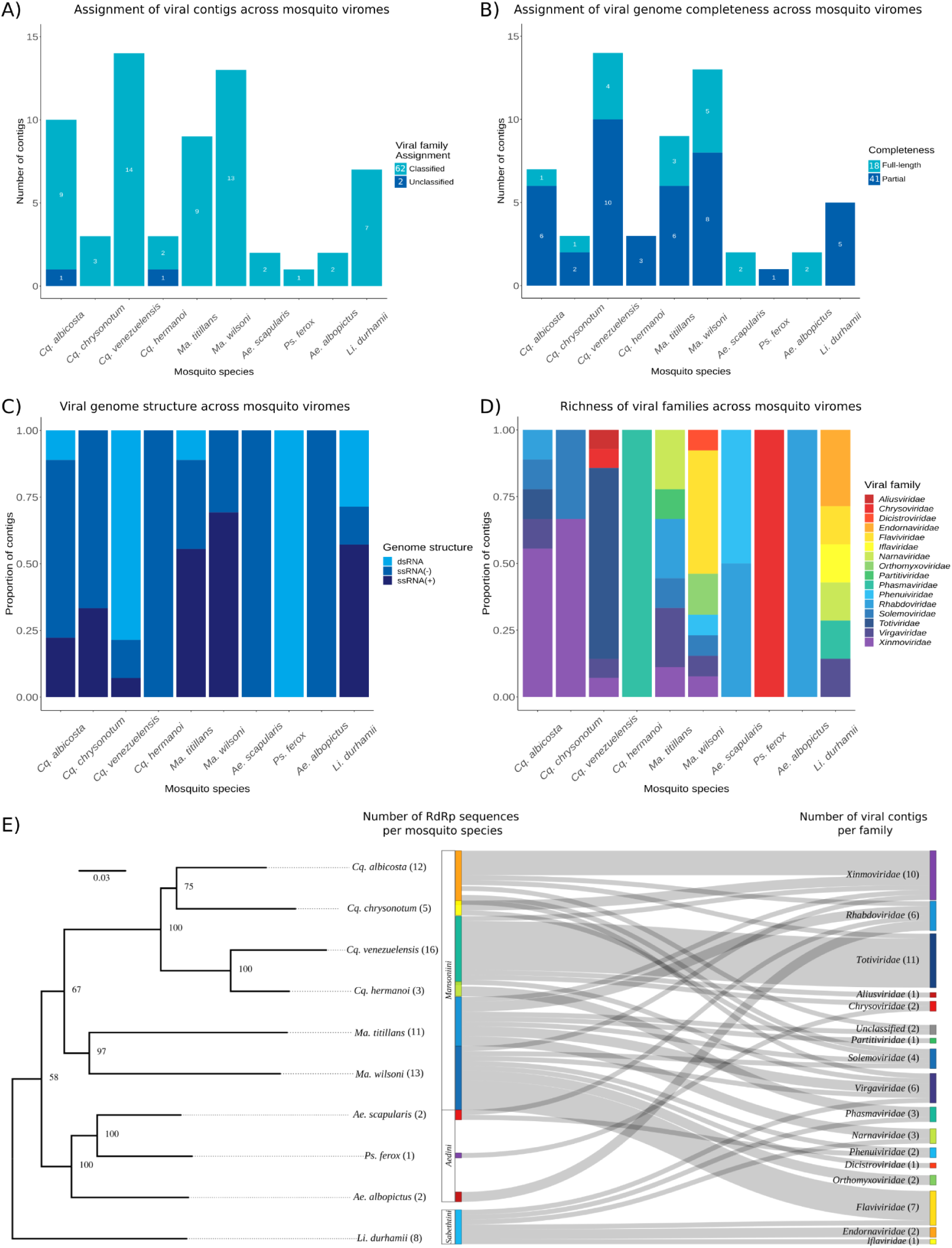
Virome composition across studied mosquito species. A) Stacked plot presenting the number of viral contigs assigned or not assigned to known viral families across mosquito viromes. B) Stacked plot showing the viral genome completeness of identified viral contigs. Number of viral sequences assigned as Full-length or partial from this study analyzed by the ViralComplete tool. C) Stacked plot showing the different viral genome structures of identified viral contigs across the mosquito viromes. Genome structure information for each family was obtained from the NCBI Virus database. D) Stacked plot displaying the proportion of viral families identified among mosquitoes. E) Phylogenetic tree of mosquito species with a sankey diagram of viral families identified across mosquitoes. The phylogenetic tree was reconstructed based on the mitogenomes assembled from sequenced data using MITObim 1.9 and analyzed on IQ-TREE 2.0 performing the ultrafast bootstraping with 1,000 replicates and the evolutionary model GTR+F+G4 selected by the ModelFinder. The root tree was set in an outgroup (*Drosophila melanogaster* - U37541.1 that was pruned from the tree).

Regarding the mosquito species, *Cq. venezuelensis, Ma. wilsoni* and *Li. Durhamii* showed the highest viral contig content, respectively (**Table 1**). While, *Ps. ferox* and *Ae. scapularis* showed the smallest number (**Table 1**). The viral contig sequences showed a wide length distribution ranging from 602 bp up to 13,088 kb with a median value of 2,832 bp (**Supplementary material 1 - Figure 6**). We were able to recover 5 full-length viral sequences for *Ma. wilsoni* assigned in *Xinmoviridae* (N=1), *Orthomyxoviridae* (N=2), *Virgaviridae* (N=1) and *Phenuiviridae* (N=1) families, while only one full-length sequence for each *Cq. albicosta* and *Cq, chrysonotum* species, representing viruses from *Rhabdoviridae* and *Solemoviridae* respectively (**Table 1, Supplementary material 1 - Figure 4** and **Supplementary file 1**). Two mosquito species (*Ps. ferox* and *Cq. hermanoi*) did not show any full-length sequences.

The virome profile of the mosquito species investigated here have shown a similar pattern of viral families across the mosquito tribes (**Figure 2D-E** and **Supplementary material 1 - Figure 7**). The virome of *Limatus durhamii* from the *Sabethini* tribe have shown the most distinct profile (**Figure 2D**). In the *Aedini* tribe (*Aedes* and *Psorophora* genera) we identified a limited number of viral contigs that allowed us to have only a narrow view of the virome from these species. While the *Coquillettidia and Mansonia* species from *Mansoniini* have shown a similar viral family profile among them (**Figure 2D-E**).

In the *Li. durhamii* species, we identified six viral families: *Iflaviridae, Phasmaviridae, Flaviviridae, Narnaviridae, Virgaviridae and Endornaviridae*. Two families were exclusively identified in this mosquito species (*Iflaviridae* and *Enfdornaviridae*). Within the *Aedini* species, a few contigs were classified into *Rhabdoviridae* family and identified in both *Ae. albopictus* and *Ae. scapularis. Phenuiviridae* was only identified on the *Ae. scapularis* pool and *Chrysoviridae* from *Ps. ferox* (**Figure 2D-E** and **Supplementary material 1 - Figure 7**). Excluding the *Chrysoviridae* family which infects predominantly plants and fungi, the viral families identified in *Ae. albopictus* and *Ae. scapularis* are known to infect both arthropods and vertebrates (**Figure 2E** and **Supplementary material 1 - Figure 7**).

In the *Mansoniini* viromes, both species from *Mansonia* and *Coquillettidia* genera were infected by viruses belonging to the viral families *Xinmoviridae, Virgaviridae* and *Solemoviridae* (**Figure 2D-E** and **Supplementary material 1 - Figure 7**). However, several other viral families were uniquely identified in specific species within *Mansoniini* such as *Chrysoviridae* and *Aliusviridae* in *Cq. venezuelensis, Phasmaviridae* in *Cq. hermanoi, Narnaviridae* and *Partitiviridae* in *Ma. titillans, Orthomyxoviridae, Flaviviridae, Phenuiviridae* and *Dicistroviridae* in *Ma. wilsoni.* In general *Mansoniini* mosquitoes showed viral sequences belonging to viral families known to infect arthropods, vertebrates, birds, fungi and plants (**Supplementary material 1 - Figure 7**). The *Ma. wilsoni* species revealed to have the highest number (7) of viral families from *Mansonini* species studied here, showing the following viral families: *Dicistroviridae, Xinmoviridae, Flaviviridae, Phenuiviridae, Orthomyxoviridae, Solemoviridae* and *Virgaviridae*. While only *Phasmaviridae* was identified in *Cq. hermanoi*.

### Phylogenetic reconstruction of virus relationships history

Based on the 16 viral families identified in the mosquito viromes we reconstructed a total of 15 phylogenetic trees using the RdRp hallmark gene (**Supplementary material 1 - Figure 8-22**) representing 14 viral families (*Aliusviridae, Chrysoviridae, Flaviviridae, Iflaviridae, Narnaviridae, Orthomyxoviridae, Partitiviridae, Phasmaviridae, Phenuiviridae, Rhabdoviridae, Solemoviridae, Totiviridae, Virgaviridae* and *Xinmoviridae*). Besides those, we also reconstructed a phylogenetic tree for one viral sequence with unclassified assignment. From now, we will present the results of viral families known to infect arthropods (*Aliusviridae, Xinmoviridae, Iflaviridae* and *Phasmaviridae*) and those with dual host infection behavior, infecting arthropods and vertebrates (*Flaviviridae, Phenuiviridae, Orthomyxoviridae* and *Rhabdoviridae*) (**Figure 3** and **Figure 4**). Phylogenetic trees of viral families with known plants, fungi, protozoa and oomycetes hosts that were also identified from the mosquito viromes can be found in **Supplementary material 1 (Figure 16-21)**.

**Figure 3.**
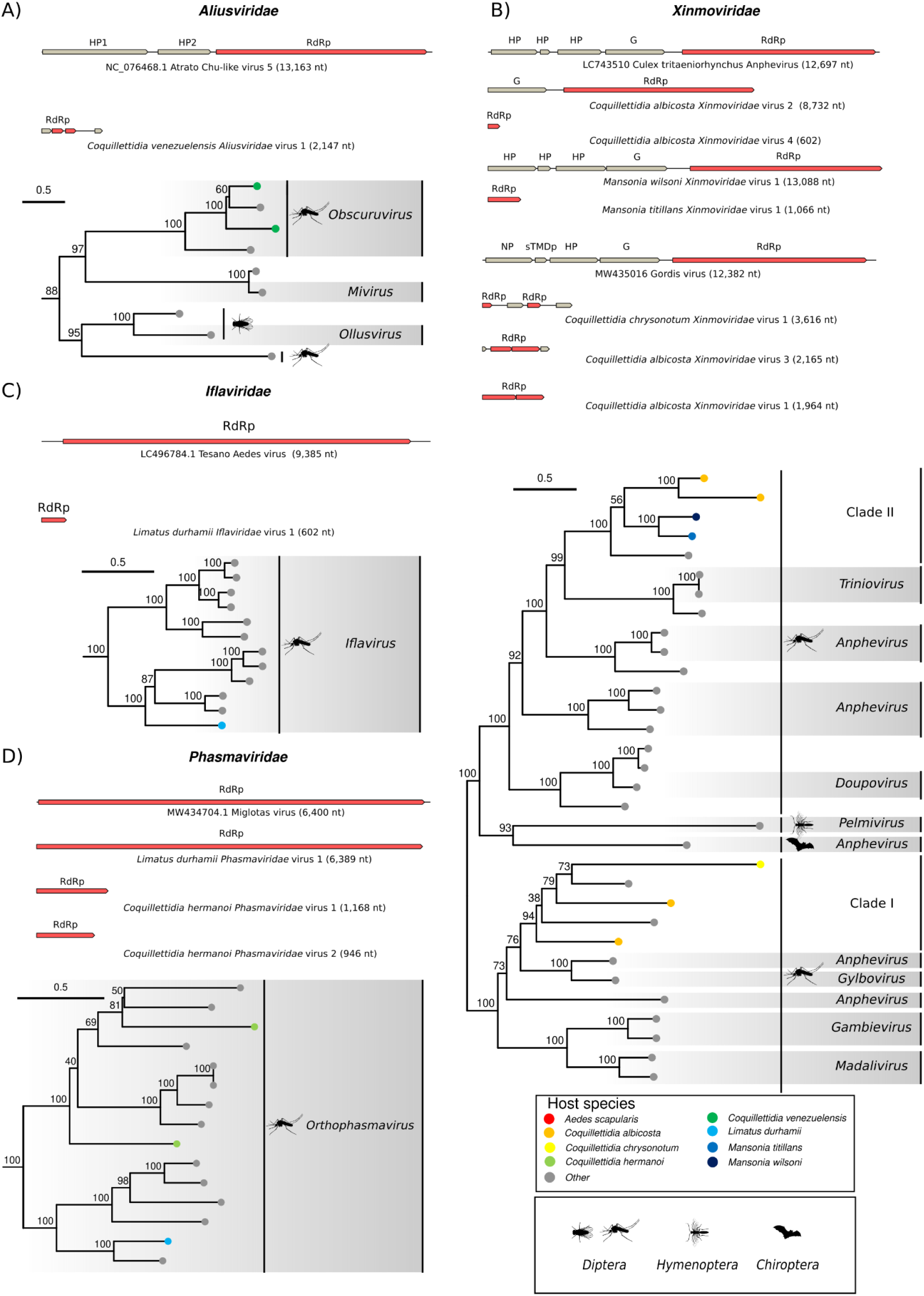
Phylogenetic trees of the *Aliusviridae*, *Xinmoviridae, Iflaviridae* and *Phasmaviridae* families. The phylogenetic trees were reconstructed based on aligned sequences representing the RdRp identified for each viral family and analyzed on IQ-TREE2.0 performing the ultrafast bootstraping with 1,000 replicates. Full phylogenetic trees can be seen in **Supplementary material 1 - Figure 8-11**. The trees were set as the midpoint root. The tip colors represent the different mosquito host species from this study, while silhouettes represent the host species recovered from the NCBI. Gray boxes indicate the subgenus of viruses according to information from NCBI and ICTV.

**Figure 4.**
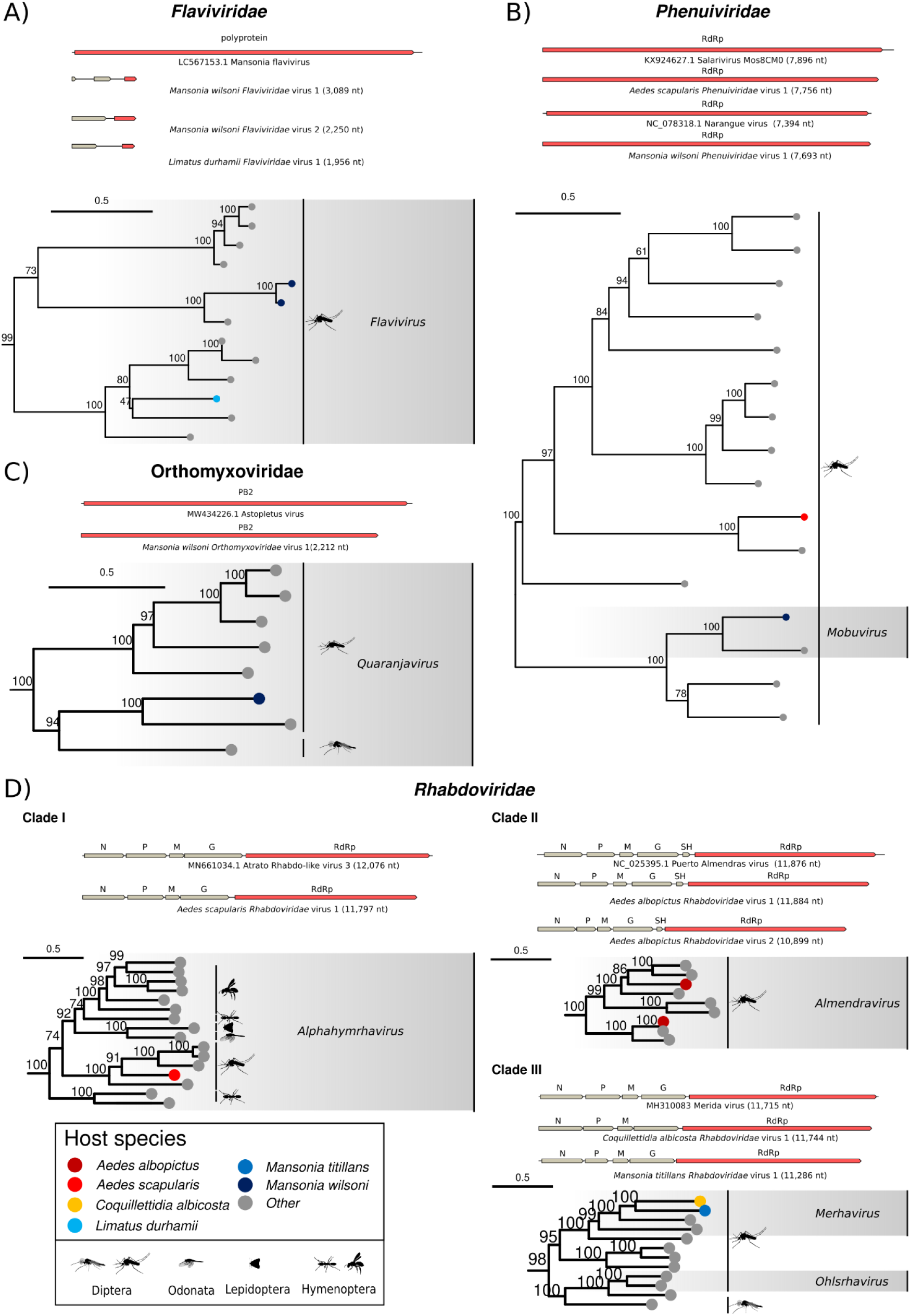
Phylogenetic trees of *Flaviviridae, Phenuiviridae, Orthomyxoviridae* and *Rhabdoviridae* families. The phylogenetic trees were reconstructed based on aligned sequences representing or the RdRp or polyprotein sequences for *Flaviviridae* and PB2 for *Orthomyxoviridae* family and analyzed on IQ-TREE2.0 performing the ultrafast bootstrapping with 1,000 replicates. Full phylogenetic trees can be seen in **Supplementary material 1 - Figure 12-15.** The trees were set as the midpoint root. The tip colors represent the different mosquito host species from this study, while the silhouettes represent the host species recovered from the NCBI. Gray boxes indicate the subgenus of viruses according to information from NCBI and ICTV. Red regions from the genomic map represent the sequence regions used in this analysis.

### Arthropod exclusive viruses

#### Aliusviridae family

Two viral sequences encoded from a contig (*Coquillettidia venezuelensis Aliusviridae virus 1*) identified in *Cq. venezuelensis* were analysed (**Figure 3A** and **Supplementary material 1 - Figure 8**). All sequences analyzed from *Coquillettidia venezuelensis Aliusviridae virus* 1 grouped into a clade together with Atrato Chu−like virus 5 (YP_010798469.1), a virus originally identified in *Ps. albipes*. These viral sequences showed an RdRp identity between 56% to 62% (**Supplementary material 1 - Figure 23** and **Supplementary file 2**). The *Coquillettidia venezuelensis Aliusviridae virus* 1 was classified as members of *Obscurusvirus* according to phylogenetic positioning and grouped in a major clade with other mosquitoes derived sequences.

#### Xinmoviridae family

For the *Xinmoviridae* family we reconstructed a tree including seven sequences identified in *Cq. albicosta* (N=4)*, Cq. chrysonotum* (N=1) and *Ma. wilsoni* (N=2) (**Figure 3B** and **Supplementary material 1 - Figure 9)**. In general our sequences were placed into two distinct clades (**Figure 3B** and **Supplementary material 1 - Figure 9**). The clade I grouping sequences from *Cq. albicosta* (*Coquillettidia albicosta Xinmoviridae virus 3* and *Coquillettidia albicosta Xinmoviridae virus 1*) and *Cq. chrysonotum* (*Coquillettidia chrysonotum Xinmoviridae virus 1*) that grouped together Gordis virus (QRW42745.1) and Culex mononega−like virus 1 (QGA70931.1). The sequence from *Coquillettidia albicosta Xinmoviridae virus 1* showed to be more divergent being placed as a sister branch of the remaining sequences of *Cq. albicosta* and *Cq. chrysonotum* (**Supplementary material 1 - Figure 9**). However the positioning inside the clade was not supported due the low UFboot value for most of the nodes.

The clade II grouped sequences from *Cq. albicosta* (*Coquillettidia albicosta Xinmoviridae* virus 2) and viral sequences from *Ma. wilsoni* (*Mansonia wilsoni Xinmoviridae virus 1*), while the basal sequence from this clade was Malby virus (UYE93836.1), a virus identified in *Aedes communis* from Sweden (**Supplementary material 1 - Figure 9**). A distinct separation of viruses from *Mansonia* and *Coquillettidia* was evident. Specifically, *Mansonia wilsoni Xinmoviridae virus 1* and *Mansonia titillans Xinmoviridae virus 1* that clustered together in a subclade. While *Coquillettidia albicosta Xinmoviridae* virus 2 and *Coquillettidia albicosta Xinmoviridae* virus 4 were grouped together into a separated subclade (**Supplementary material 1 - Figure 9**). The identity values of our sequences with Malby virus ranged from 47% to 59% suggesting a high divergence of those sequences in relation with sequences from the literature (**Supplementary material 1 - Figure 23** and **Supplementary file 2**). In general, both major clades were represented by sequences derived from mosquito hosts.

#### Iflaviridae

For the *Iflaviridae* family we identified one sequence derived from a virus that we designated as *Limatus durhamii Iflaviridae virus 1* in *Li. durhamii* species. Our phylogenetic tree revealed this virus grouping together other viruses from *Iflavirus* genus (**Figure 3C** and **Supplementary material 1 - Figure 10**). However, *Limatus durhamii Iflaviridae* virus 1 has shown to be highly divergent from viruses in the literature. This virus was placed as a basal branch together with other viruses identified mainly in *Aedes* and *Culex spp.* The RdRp identity values among *Limatus durhamii Iflaviridae* virus 1 and those viruses inside this clade ranged from 55 to 59% of identity (**Supplementary material 1 - Figure 23** and **Supplementary file 2**).

#### Phasmaviridae family

For the *Phasmaviridae* family we positioned three sequences from *Cq. hermanoi* (N=2) and *Li. durhamii* (N=1) (**Figure 3D** and **Supplementary material 1 - Figure 11**). Our sequences were assigned into *orthophasmavirus* genus and clustered in distinct clades. The sequence from *Limatus durhamii Phasmaviridae* virus 1 was placed together with Miglotas virus (QRW41773.1), identified in *Culex erythrothorax* from the United States. *Coquillettidia hermanoi Phasmaviridae* virus 1 is more divergent and was placed as a sister branch of other *Phasmaviridae* viruses identified in *Culex, Aedes* and *Mansonia* species, besides the *Coquillettidia hermanoi Phasmaviridae* virus 2 (**Supplementary material 1 - Figure 11**). The RdRp identity among *Coquillettidia hermanoi Phasmaviridae* virus 1 and other viruses from the clade ranged from 31 to 63%.

### Dual host viruses infecting arthropod and vertebrates

#### Flaviviridae family

For the *Flaviviridae* tree we analyzed three sequences that were placed into *Flavivirus* genus and we defined as *Mansonia wilsoni Flaviviridae* virus 1*, Mansonia wilsoni Flaviviridae* virus 2 and *Limatus durhamii Flaviviridae* virus 1 (**Figure 4A** and **Supplementary material 1 - Figure 12**). The sequences from *Mansonia* and *Limatus* were placed into distinct clades, where the viruses derived from *Ma. wilsoni* clustered together with *Mansonia* flavivirus (BCI56826.1) that was characterized from mosquitoes of the *Mansonia* genus in Bolívia. The identities of analyzed sequences ranged from 65 to 72% that suggest a high divergence of the Flaviviridae viruses identified here in relation to those available in the literature. The *Limatus durhamii Flaviviridae* virus 1 was placed into another clade together with other viruses identified in *Sabethes, Culex* and *Culiseta* mosquito genera. However their positioning inside the *Flaviviridae* tree was not solved due the low UFBoot values for node support.

#### Phenuiviridae family

For the *Phenuiviridae* family we positioned the two sequences representing complete segments identified in *Ae. scapularis* and *Ma. wilsoni* (**Figure 4B** and **Supplementary material 1 - Figure 13**). Our *Phenuiviridae* viral sequences were grouped in two distinct clades. *Mansonia wilsoni Phenuiviridae* virus 1 grouped together with the Narangue virus (YP_010839995.1) with an RdRp identity of 65%. This virus was characterized from *Ma. titillans* species in Colombia. While the *Phenuiviridae* sequence from *Ae. scapularis* was placed together with Salarivirus Mos8CM0 (API61884.1) showing an RdRp amino acid identity of 65%. This virus was identified firstly in non-identified mosquito samples from the United States.

#### Orthomyxoviridae family

For the *Orthomyxoviridae* family we reconstructed a tree comprising one sequence of PB2 subunit that compose the RdRp of *Orthomyxoviridae* viruses (**Figure 4C** and **Supplementary material 1 - Figure 14**). The sequence derived from *Mansonia wilsoni Orthomyxoviridae* virus 1 grouped into a clade of sequences classified as *Quaranjavirus* and was close to Astopletus virus (QRW42566.1) showing high divergence with a PB2 amino acid identity of 46% (**Figure 4C** and **Supplementary material 1 - Figure 14**).

#### Rhabdoviridae family

For the *Rhabdoviridae* family we included five sequences representing complete genomes detected in *Ae. scapularis* (N= 1)*, Ae. albopictus* (N=2)*, Ma. titillans* (N=1) and *Cq. albicosta* (N=1) (**Figure 3D** and **Supplementary material 1 - Figure 15**). In general our sequences were clustered into 3 distinct clades and classified as members of *Alphahymhavirus* (*Aedes scapularis Rhabdoviridae* virus 1), *Almendravirus* (*Aedes albopictus Rhabdoviridae* virus 1 and *Aedes albopictus Rhabdoviridae* virus 2) and *Merhavirus* (*Mansonia titillans Rhabdoviridae* virus 1 and *Coquillettidia albicosta Rhabdoviridae* virus 1) genera, respectively. The *Aedes scapularis Rhabdoviridae* virus 1 grouped into a clade together other viruses identified in *Aedes* and *Culex* genera such as Atrato Rhabdo−like virus 3 (QHA33680.1), San Gabriel mononegavirus (DAZ85658.1), Primus virus (QIS62334.1) and XiangYun mono−chu−like virus 4 (UUG74104.1). The *Aedes scapularis Rhabdoviridae* virus 1 was positioned as a basal branch in relation to San Gabriel mononegavirus, Primus virus and XiangYun mono−chu−like virus 4 (**Figure 4D - Clade I** and **Supplementary material 1 - Figure 15**) and showed an RdRp amino acid identity of 52% with Atrato Rhabdo−like virus 3. The two viruses identitified in *Ae. albopictus* (*Aedes albopictus Rhabdoviridae* virus 1 and *Aedes albopictus Rhabdoviridae* virus 2) were positioned in two distinct branches within the *Almendravirus* clade (**Figure 4D - Clade II**). Specifically, *Aedes albopictus Rhabdoviridae* virus 1 clustered closely with Puerto Almendras virus (YP_009094394.1), a virus identified in *Aedes fulvus* from Peru. Our analysis revealed that these two viruses are probably the same virus given the remarkably high amino acid identity of the RdRp - 98%. The other *Almendravirus* identified in *Ae. albopictus* was *Aedes albopictus Rhabdoviridae* virus 2 that clustered together other *Almendraviruses* identified in *Aedini* mosquitoes from *Psorophora* and *Armigeres* genera (**Figure 4D - Clade 2** and **Supplementary material 1 - Figure 15**). The last clade of *Rhabdoviridae* sequences identified in *Mansoniini* species from this study, clustered *Mansonia titillans Rhabdoviridae* virus 1 and *Coquillettidia albicosta Rhabdoviridae* virus 1 together with Merida virus (AWJ96718.1), a virus identified in *Cx. quinquefasciatus* from the United States. The identity of RdRp from those viruses with Merida virus was approximately 49%.

## Discussion

Mosquitoes carry an abundant and diverse viral community. A recent review has shown that viruses were investigated and detected in at least 128 mosquito species and 14 genera around the globe (Moonen et al., 2023). Most of theses viruses have been identified in species of *Culex, Aedes* and *Anopheles* genera, that have been the focus of extensive research efforts, once they are medically important vectors (Atoni et al., 2019b; de Almeida et al., 2021; Moonen et al., 2023; Parry et al., 2021). However, it is crucial to study the virome composition from other mosquito genera and species, mainly those from sylvatic and urban-sylvatic interfaces once the majority of arthropod-borne viruses have emerged and are maintained in the sylvatic environment and may be transmitted to humans through bridge vectors (Weaver, 2005). Studying the virome from sylvatic mosquito could not only bring information from novel viruses that are maintained in nature, but also pinpoint potentially human pathogenic viruses as well as reveal non-pathogenic viruses that could have potential to reduce replication from other arboviruses.

While metatranscriptome sequencing offers an unbiased approach to characterizing a diverse number of viruses, it presents some challenges for accurate viral identification. Previous studies have highlighted that some factors, including the choice of assembler can impact on virome characterization (Roux et al., 2017; Sutton et al., 2019). Additionally, the use of specific databases on similarity sequence methods can also impact viral identification. With the aim of reducing the knowledge gap in virome of sylvatic mosquitoes from Brazil, we conducted metatranscriptome sequencing of 10 distinct species and evaluated different viral identification approaches coupled with different viral databases. The databases included in this study encompassed sequences available on NCBI to specialized RdRp databases that included divergent and unclassified viral sequences that are not present in most popular databases. Our analysis indicates that conventional databases containing viral proteins are generally adequate for detecting the majority of viruses including more divergent viruses. However, some specific viral contigs were exclusively identified using specialized databases or using HMM models. Therefore, coupling of different approaches and databases on viral identification offer a comprehensive and complementary understanding of virome composition.

Our analysis identified a total of 64 viral sequences bearing RdRp domain representing 16 viral families. Then, we could reassess the virome composition of those two mosquito species from a previous study (*Ma. wilsoni* and *Cq. hermanoi* from da Silva et al., 2021) and we expanded the knowledge including another eight sylvatic species from five genera, some those screened for viruses for the first time. Our findings revealed a diverse set of viruses that are highly divergent from known viruses from databases and are likely insect specific viruses within the virome of studied mosquitoes. With exception of two sequences that showed high amino acid RdRp identity (> 90%) with viral sequences available in the literature, probably representing the same virus. The majority of RdRp sequences were distinct across mosquito species, genera and from those already available in the databases (**Supplementary material 1 - Figure 23** and **Supplementary material file 2**).

In Brazil there have been few studies focusing on the virome of mosquitoes and the ones publishes covered a limited number of mosquito species from different biomes such as Cerrado, Caatinga, Pantanal, Amazon and Atlantic forest from Southeast and Northeast Brazil (Aragão et al., 2023; da Silva et al., 2021a; da Silva Ferreira et al., 2020b; da Silva Neves et al., 2021; Maia et al., 2019b; Pinto et al., 2017; Scarpassa et al., 2019b).

### Virome of *Sabethini* tribe

Regarding *Sabethini* viromes, few studies have been conducted only on species of the *Sabethes* and *Wyeomyia* genera (Aragão et al., 2023; Maia et al., 2019b; Pinto et al., 2017). Maia et al. (2019) identified viruses from *Flaviviridae, Chuviridae, Reoviridae, Phenuiviridae* and *Partitiviridae* from *Sabethes gymonothorax* salivary glands sampled in at high Pantanal, Central-Western Brazil. Another study conducted by Aragão et al. (2023) identified viral sequences from *Virgaviridae* in *Sa. chloropterus*, *Lispiviridae, Partitiviridae* and *Parvoviridae* in *Sa. quasicyaneus* and *Xinmoviridae* in *Sa. glaucodaemon*, respectively. While the two studies investigating the virome of the *Wyeomyia* genus samples in Brazil were not able to find viral sequences (Maia et al., 2019b; Pinto et al., 2017). The current study represents the first analysis of a virome of a mosquito from the *Limatus* genus in Brazil revealing the presence of six viral families (*Iflaviridae, Flaviviridae, Narnaviridae, Virgaviridae, Phasmaviridae* and *Endornaviridae*) in which *Endornaviridae* family was the most abundant and *Virgaviridae* the most abundant (**Supplementary material 1 - Figure 8**). Although human pathogenic *Flaviviridae* sequences were identified in this mosquito species (Barrio-Nuevo et al., 2020), the phylogenetically studied sequence from this viral family grouped with other ISVs (**Supplementary material 1 - Figure 12**).

### Virome of *Aedini* tribe

Regarding *Aedini* mosquitoes, some studies have assessed the virome of *Ae. albopictus* and *Ae. aegypti* (He et al., 2021), *Psorophora* species (Charles et al., 2018), *Ochlerotatus* (Truong Nguyen et al., 2022) and *Armigeres* (Thongsripong et al., 2021). Here we analyzed the virome of three *Aedini* species, our results showed a low diversity of viral families within pools analyzed (**Figure 2**). The *Ae. scapularis* showed the highest diversity within the tribe, but with only two viral families, while both *Ae. albopictus* and *Ps. ferox* showed only one viral family each. The *Aedini* species showed the lowest diversity of viral families in relation to other genera also assessed in the current study such as *Limatus, Mansonia* and *Coquillettidia* mosquitoes (**Figure 2D-E**). Another study has also shown the virome of *Ae. albopictus* with lower number of viral sequences in relation to *Ae. aegypti* mosquitoes in Colombia (Calle-Tobón et al., 2022). On the other hand, a study on the virome of *Ae. albopictus* from China revealed a higher number of identified viral families, with at least 50 families specifically associated with vertebrates, invertebrates, plants, fungi, bacteria, and protozoa hosts (He et al., 2021). In Brazil, there have been few studies investigating the virome in *Aedini*, and the outcomes are similar with our findings, where the species exhibited a lower number of viral families or in certain cases did not showing viral sequences (Aragão et al., 2023; Duarte et al., 2022; Maia et al., 2019b; Pinto et al., 2017; Ribeiro et al., 2020). Each of these studies were conducted using distinct mosquito sampling, sequencing and bioinformatics approaches hindering direct comparison of the results. More studies focusing on the virome of *Aedini* species in different geographical regions are needed to uncover the main differences in viral diversity among *Adini* species around the world.

### Virome of *Mansoniini* tribe

*Ma. wilsoni* and *Cq. hermanoi* species from the *Mansoniini* tribe were already studied from the virome perspective by our own group. Characterizing the virome of different samples of the same species allowed us to recover five viral families identified before (*Orthomyxoviridae, Xinmoviridae, Phenuiviridae, Flaviviridae* and *Virgaviridae*). Moreover, we characterized another three which were reported for the first time: *Phasmaviridae* for *Cq. hermanoi and Dicistroviridae* and *Solemoviridae* for *Ma. wilsoni*. These viral families are known to infect Arthropods, and Plants, respectively. However, we have not identified *Rhabdoviridae*, *Chuviridae* and *Partitiviridae* viral sequences in those species as previously identified (da Silva et al., 2021a). Our analysis revealed the presence of the Aliusviridae family belonging to the *Jingchuvirales* order in *Cq. venezuelensis*. Recently, the *Jingchuvirales* order originally consisted only of the Chuviridae family that was splitted into five viral families, including the *Aliusviridae* family (Di Paola et al., 2021). In general, our results for *Mansoniini* were similar as observed in a previous study (Thongsripong et al., 2021) wherein the majority of sequences from *Ma. uniformis* were classified into the *Xinmoviridae* family with exception of *Cq. hermanoi* where the majority of sequences belonged to *Phasmaviridae*. In Brazil, the only study from another research group assessing viruses from *Ma. wilsoni* has identified a few sequences classified in *Totiviridae*, *Partitiviridae* and *Chuviridae* (Pinto et al., 2017). However those authors used a different mosquito tissue (salivary glands) and sequencing approach, which may explain the differences found (Pinto et al., 2017).

### Arthropods and vertebrates viruses

We identified some viral contigs belonging to families (*Phenuiviridae, Orthomyxoviridae, Rhabdoviridae, Flaviviridae*) known to harbor viruses that infect arthropods and vertebrates (**Figure 1E**). The *Phenuiviridae* family encompasses arboviruses such as the Rift Valley Virus which is transmitted mainly by *Aedes* and *Culex* mosquitoes (Kwaśnik et al., 2021; Sun et al., 2022). In our study we positioned two *Phenuiviridae* sequences, one from *Ae. scapularis* that is clustered with Salarivirus Mos8CM0 (API61884.1) and one from *Ma. wilsoni* grouping with Narangue virus (QHA33858.1) (**Supplementary material 1 - Figure 10**). Due to the limited knowledge about these viruses further studies are required to evaluate their effect in different cell lines and their infection capacity in invertebrate and vertebrate cells.

The *Orthomyxoviridae* also encompass known arboviruses that can infect both arthropods and vertebrates such as *Quaranjarviruses* (Quaranfil virus) and *Thogotovirus* (Thogotovirus Thogoto) (Hubálek and Rudolf, 2012; Presti et al., 2009). Here we recovered the *Orthomyxovirus* previously detected in *Ma. wilsoni* (da Silva et al., 2021a). The *Ma. wilsoni Othomyxoviridae* virus 1 was assigned into the quaranjavirus genus. There is evidence of vertebrate and humans cell infection for viruses of this genus highlighting its medical significance (Allison et al., 2015; Taylor et al., 1966).

Viruses from the *Rhabdoviridae* are generalist and were identified into a wide range of hosts (Dietzgen et al., 2017). The sequences generated in this study clustered with other *Rhabdoviridae* viruses found in mosquitoes. Interestingly, one clade that clustered the sequences identified in *Ae. albopictus* revealed to be a basal clade encompassing viruses known to infect a wide range of vertebrates and arthropods (**Supplementary material 1 - Figure 14**). The sequences of this basal clade have been assigned in Almendravirus genus, which includes mosquito Rhabdoviruses. One sequence identified in *Ae. albopictus* (*Ae. albopitus Rhabdoviridae* virus 1) showed a high amino acid identity with Puerto Almendras virus that was previously identified in *Aedes fulvus* in Peru (Vasilakis et al., 2014). Although this virus cause no cytopathic effect in vertebrates cells it could be detected in the cell culture supernatant suggesting active replication in vertebrate cells (Vasilakis et al., 2014).

Our phylogenetic analysis revealed that the sequences from *Flaviviridae, Rhabdoviridae, Phenuiviridae* and *Orthomyxoviridae* clustered with other ISVs that have been previously characterized although with substantial amino acid divergence. Based on known hosts of these clades, our findings suggest we characterized several mosquito-specific viruses (MSV) (**Figure 3** and **Figure 4**). Our results are consistent with previous studies that have also found a low proportion of pathogenic arboviruses among the metatranscriptomic uncovered viruses (de Almeida et al., 2021; Moonen et al., 2023; Shi et al., 2017, 2016). The MSV capacity to infect vertebrate hosts remains unclear, and further research is needed to assess the infection potential of these novel viruses.

### Arthropod viruses

Several viral contigs identified in this study were classified within known viral families bearing members that interfere with arboviruses replication such as *Xinmoviridae* (*Anphevirus* genus), *Phasmaviridae* and *Phenuiviridae* families. ISVs from those families such as Phasi Charoen virus (PCV) from *Phenuiviridae* family together Cell fuging agent virus from the *Flaviviridae* have been shown to interfere with replication of arboviruses such as DENV, ZIKV and La Crosse Virus from *Flaviviridae* and *Peribunyaviridae* respectively (Schultz et al., 2018). Furthermore Olmo et al’s study showed a wide distribution of Phasi Charoen Like Virus on *Aedes* mosquitoes in different locations and its influence in DENV and ZIKV transmission *in vivo* (Olmo et al., 2023). No viral contig with similarity with this virus was found in our dataset. One study has described the capacity of Aedes anphevirus from the *Xinmoviridae* family to interfere with Dengue virus replication in cell lines (Parry and Asgari, 2018). We found several contigs from both *Mansonia* and *Coquillettidia* genera that were clustered in a major clade with Aedes anphevirus.

In summary, we uncovered the virome of sylvatic mosquitoes from Northeast Brazil. We detected a diverse virome largely composed of highly divergent ISVs or MSVs.

## Limitation of the study

The analysis of virome composition of eukaryotic species including mosquitoes has revealed a substantial dynamism through time linked with some specific factors that affect viral richness and abundance (Atoni et al., 2018; Pettersson et al., 2019; Shi et al., 2017; Xia et al., 2018). Factors such as sampling collection in different seasons (Feng et al., 2022; He et al., 2021) or host feeding pattern (Shi et al., 2022) have shown to influence the mosquito virome. In this study, we did not evaluate the virome composition through time and limited our sampling to a narrow species range distribution, therefore our data is a snapshot of the viral diversity carried by sylvatic mosquitoes. Moreover, due to the different mosquito tissues, viral enrichment, sequencing and bioinformatic analysis performed in other studies from Brazil and the world we could not directly compare the virome abundance and diversity with our findings. Anyhow, the data presented here is in line with a broad picture found in other studies in the sense that mosquito viromes are mainly composed of highly divergent ISVs. Therefore, a more intense sampling and virome characterization using less biased wet and dry lab protocols is needed to fully uncover sylvatic mosquitoes viromes in Brazil.

Virome findings may be biased by the presence of endogenous viral elements (EVEs), viral remnants endogenized into host genomes that can be transcribed (de Almeida et al., 2021; Nouri et al., 2018). Insects, including mosquitoes, have been shown to harbor several RNA virus EVEs into their genomes (Wallau, 2022). In order to differentiate EVEs from bonafide circulating viruses one could compare the identified viruses’ genomes against the host genomes, as performed in previous studies (da Silva et al., 2021a; Zakrzewski et al., 2018). Due to the lack of host genomic sequences and small RNA datasets for the majority of mosquitoes investigated here we compared the identified viral sequences against all genomes available for mosquito species and included draft genomes for some species studied here amining to reduce the interference of EVEs in our virome analysis. Therefore, further assessment of bonafide viruses found here is important to validate the findings. Although some of viral contigs identified warrant further in-depth analysis to discriminate them from EVEs, we identified several complete viral sequences that likely represent true bona fide viruses once the large majority of EVEs are composed of short fragments of viral genomes (Katzourakis and Gifford, 2010; Ter Horst et al., 2019).

It is also important to point out that we focused our analysis on the most conserved RdRp gene/protein once it is the best marker for detecting and performing evolutionary studies with distantly related viruses, but we did not performed detailed analysis of other viral genes, this can particularly bias the viral completeness results regarding segmented viruses. Hence, future analysis is warranted to more in depth characterize segmented virus genomes completeness. Moreover, the metatranscriptomic approach used is better suited to characterize viruses with RNA genome, but it can also detect actively transcribed DNA viruses. Therefore, we have here analyzed more specifically the sylvatic mosquito RNA virome and the viral community with DNA genomes should be further accessed.

## Conclusion

In this study we conducted a virome profiling of 10 mosquito species from Northeast Brazil and identified viral sequences from 16 different families, including a total of 18 full-length sequences of viral segments or complete linear genomes. Our phylogenetic analysis revealed that the majority of the newly characterized viruses represent new viral species based on RdRp amino acid identities. Our study provides important basic knowledge about the virome composition of mosquitoes in Northeast Brazil and highlights several viral families that require further investigation to evaluate their potential to infect vertebrates or impact arbovirus replication. These findings enhanced our understanding of how these viruses are maintained in nature, their interactions with the sylvatic mosquito fauna and the natural viral community that infect those species.

## Acknowledgments

This project was supported by the Inova Fiocruz Funding program (VPPCB-007-FIO-18 and CNPq/Instituto Aggeu Magalhães - FIOCRUZ N° 39/2018). G.L.W. was supported by the Conselho Nacional de Desenvolvimento Científico e Tecnológico (CNPq) through their productivity research fellowships (303902/2019-1). Alexandre Freitas da Silva and Laís Ceschini Machado received scholarship supported by Fundação de Amparo à Ciência e Tecnologia do Estado de Pernambuco (FACEPE). Luisa Maria Inácio da Silva was supported by a scholarship from the Instituto Aggeu Magalhães (IAM) - FIOCRUZ. Filipe Zimmer Dezordi received a scholarship from Coordenação de Aperfeiçoamento de Pessoal de Nível Superior (CAPES). We thank Anna Christina de Matos Salim from Instituto René Rachou (IRR) - Fundação Oswaldo Cruz for the metatranscriptome sequencing support. We thank the Núcleo de Bioinformática of IAM and Rede de Plataformas Tecnológicas (FIOCRUZ) for the computational infrastructure and sequencing platform, respectively. We thank the Jardim Botânico do Recife and Parque Estadual Dois Irmãos for the authorization to perform the mosquito sampling.

## Author Contributions

Alexandre Freitas da Silva performed mosquito sampling, mosquito identification, data analysis and wrote the original draft. Laís Ceschini Machado performed the mosquito sampling, molecular experiments, contributed to drafting and review of the manuscript. Filipe Zimmer Dezordi performed the mosquito sampling, supported the sequencing data analysis and contributed to drafting and review of the manuscript. Luisa Maria Inácio da Silva performed the mosquito sampling, mosquito identification, molecular experiments and contributed to drafting and review of the manuscript. Gabriel Luz Wallau conceptualized, supervised and administered the project, designed the methodology and wrote, reviewed and edited the manuscript.

## Data Availability

Raw reads generated in this study are available through the European Bioinformatic Institute under the project number: PRJEB63303. Supplementary material 1 and Supplementary files 1 and 2 are available in Figshare (https://doi.org/10.6084/m9.figshare.23584827.v1).

