## Supplementary Material 1 for "Highly divergent and diverse viral community infecting sylvatic mosquitoes from Northeast Brazil"

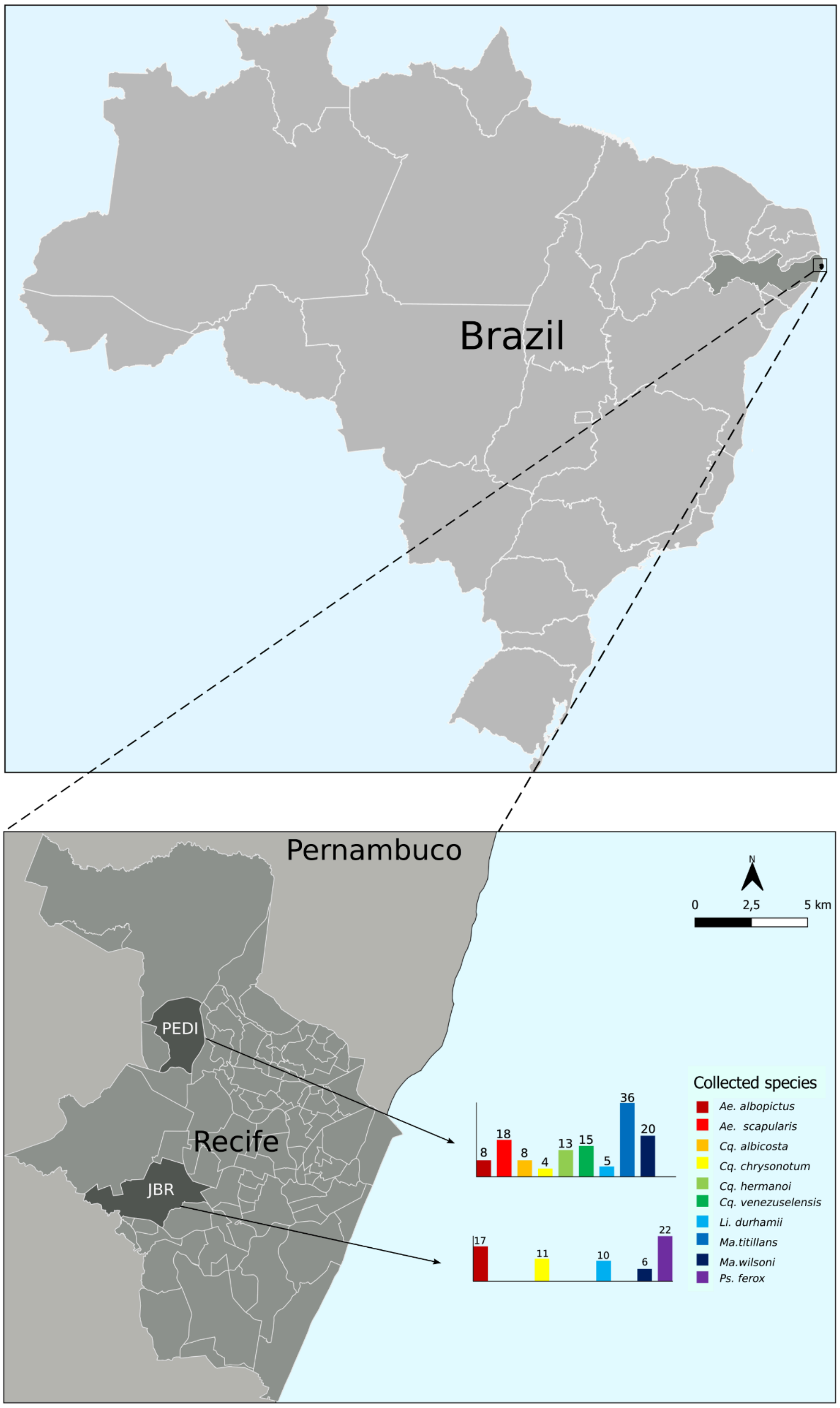


**Figure 1.** Mosquito collection points. The figure shows the map of collection points used to sampling sylvatic mosquito species from remnants of Atlantic forest in Recife municipality, state of Pernambuco, Brazil. PEDI refers to Parque Estadual Dois Irmãos (8º00′43.3″S, 34º56′40.7″W) and JBR - Jardim Botânico do Recife (8°04'33.0"S, 34°57'35.9"W).

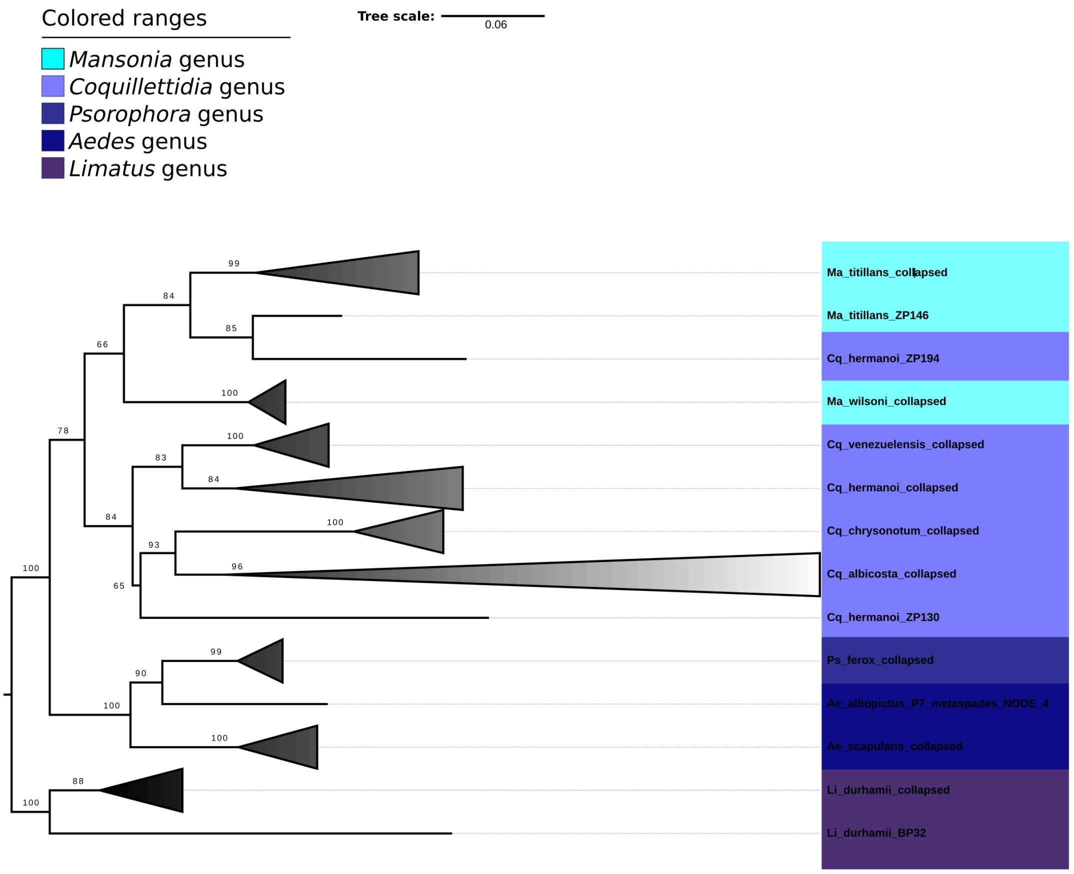


**Figure 2.** Phylogenetic tree of Cytochrome c oxidase subunit 1 (COI) from the mosquito pools. The analysis was based on the COI alignment of 605 nucleotide sites for each specimen used in this work. The tree was reconstructed using the IQ-TREE 2.0 using the evolutionary model TPM3+F+G4 and performing the ultrafast bootstrapping with 1,000 replicates. The tree was plotted on Figtree v1.4.4.

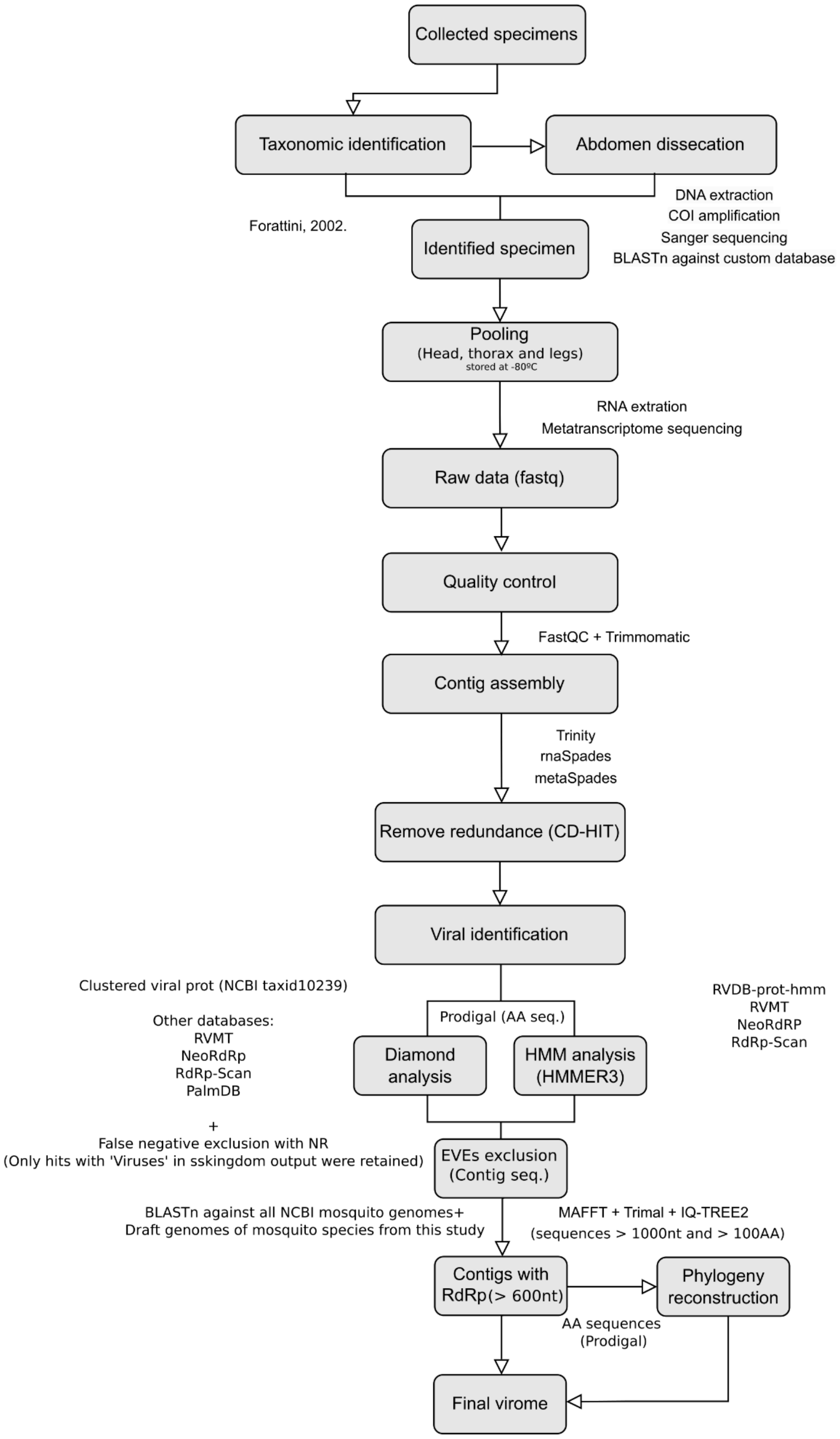


**Figure 3.** Study workflow design. Scheme presenting the steps followed from collection of mosquitoes, sequencing, bioinformatic analysis and virome identification.

**Table 1.** Terms used for RdRp filtering for NCBI and RVDB databases.

| Taxon | NCBI | RVDB-prot-HMM |
|---|---|---|
| Generalist | RNA-dependent, RNA polymerase, RdRp, L protein | RNA, polymerase |
| Virgaviridae | polyprotein | - |
| Endornaviridae | polyprotein | - |
| Flaviviridae | polyprotein, NS5 protein, NS5 and non structural protein 5 | - |
| Dicistroviridae | ORF1, nonstructural, polyprotein | - |
| Chuviridae | reverse transcriptase | - |
| Partitiviridae | reverse transcriptase | - |
| Orthomyxoviridae | PB1, PB2, PA | - |
| Luteoviridae | P1-P2 fusion protein | - |

**Table 2.** General information of NCBI mosquito genomes used on EVEs removal step.

| Organism Scientific Name | Assembly Name | Assembly Accession | Level | Contig N50 | Size | Gene Count |
|---|---|---|---|---|---|---|
| *Anopheles gambiae str. PEST* | AgamP3 | GCF_000005575.2 | Chromosome | 85548 | 265011681 | 13271 |
| *Aedes aegypti* | AaegL5.0 | GCF_002204515.2 | Chromosome | 11758062 | 1278715314 | 19623 |
| *Aedes albopictus* | Aalbo_primary.1 | GCF_006496715.2 | Scaffold | 1185651 | 2535618872 | 39747 |
| *Anopheles stephensi* | UCI_ANSTEP_V1.0 | GCF_013141755.1 | Chromosome | 38117870 | 243460342 | 15394 |
| *Anopheles albimanus* | VT_AalbS3_pri_1.0 | GCF_013758885.1 | Chromosome | 25000043 | 172602732 | 12945 |
| *Culex quinquefasciatus* | VPISU_Cqui_1.0_pri_paternal | GCF_015732765.1 | Chromosome | 2875282 | 573214445 | 18458 |
| *Culex pipiens pallens* | TS_CPP_V2 | GCF_016801865.2 | Chromosome | 794805 | 566339288 | 19673 |
| *Anopheles arabiensis* | AaraD3 | GCF_016920715.1 | Chromosome | 23935993 | 256807969 | 15763 |
| *Anopheles merus* | AmerM5.1 | GCF_017562075.2 | Chromosome | 2729089 | 294364394 | 16199 |
| *Toxorhynchites rutilus septentrionalis* | ASM2978413v1 | GCF_029784135.1 | Chromosome | 3797918 | 903034419 | 21469 |
| *Uranotaenia lowii* | ASM2978415v1 | GCF_029784155.1 | Chromosome | 600000 | 1077644186 | 23533 |
| *Wyeomyia smithii* | ASM2978416v1 | GCF_029784165.1 | Chromosome | 2828848 | 769230131 | 17568 |
| *Malaya genurostris* | Malgen_1.1 | GCF_030247185.1 | Chromosome | 7566576 | 831262279 | 15668 |
| *Topomyia yanbarensis* | ASM3024719v1 | GCF_030247195.1 | Chromosome | 2783074 | 1168159140 | 20654 |
| *Anopheles cruzii* | idAnoCruzAS_RS32_06 | GCF_943734635.1 | Chromosome | 343635 | 184069349 | 12401 |
| *Sabethes cyaneus* | idSabCyanKW18_F2 | GCF_943734655.1 | Chromosome | 22692164 | 676043992 | 14156 |
| *Anopheles aquasalis* | idAnoAquaMG_Q_19 | GCF_943734665.1 | Chromosome | 38496358 | 176573000 | 12989 |
| *Anopheles coluzzii* | AcolN3 | GCF_943734685.1 | Chromosome | 18767309 | 262602229 | 14579 |
| *Anopheles* | idAnoMacuD | GCF_9437346 | Chromosome | 22618715 | 224059441 | 12929 |

| | | | | | | |
|---|---|---|---|---|---|---|
| *maculipalpis* | A_375_x | 95.1 | | | | |
| *Anopheles coustani* | idAnoCousDA _361_x.2 | GCF_9437347 05.1 | Chromosome | 27992483 | 269983652 | 14493 |
| *Anopheles marshallii* | idAnoMarsDA _429_01 | GCF_9437347 25.1 | Chromosome | 32607359 | 225712104 | 13358 |
| *Anopheles darlingi* | idAnoDarlMG _H_01 | GCF_9437347 45.1 | Chromosome | 19203082 | 181637336 | 12393 |
| *Anopheles moucheti* | idAnoMoucS N_F20_07 | GCF_9437347 55.1 | Chromosome | 4743928 | 271316723 | 14232 |
| *Anopheles ziemanni* | idAnoZiCoDA _A2_x.2 | GCF_9437347 65.1 | Chromosome | 27992483 | 269983652 | 14085 |
| *Anopheles funestus* | idAnoFuneDA -416_04 | GCF_9437348 45.2 | Chromosome | 24104831 | 250698078 | 14819 |
| *Anopheles bellator* | idAnoBellAS_ SP24_06.2 | GCF_9437357 45.2 | Chromosome | 196687 | 169575294 | 11682 |
| *Anopheles nili* | idAnoNiliSN_ F5_01 | GCF_9437379 25.1 | Chromosome | 37444657 | 195220701 | 12354 |

**Table 3.** Draft genome assemblies from this study used for EVEs filtering. Genomic statistics generated using the gt seqstat tool.

| **Mosquito species** | **Assembly method** | **Lib. Name** | **N of contigs** | **Total contigs length** | **median contig size** | **longest contig** | **N50** |
|---|---|---|---|---|---|---|---|
| *Ma. titillans* | Megahit | E | 1,296,691 | 1,299,522,734 | 692 | 95,249 | 1,272 |
| *Ma. wilsoni* | Megahit | D+Zpp1 | 12,483,356 | 6,160,630.966 | 320 | 27,220 | 546 |
| *Ae. albopictus* | Megahit | F | 3,071,614 | 1,998,281,880 | 474 | 57,926 | 762 |
| *Ps. ferox* | - | - | - | - | - | - | - |
| *Ae. scapularis* | - | - | - | - | - | - | - |
| *Cq. chrysonotum* | Megahit | A | 3,997,033 | 2,277,546,645 | 421 | 106,288 | 622 |
| *Cq. venezuelensis* | Megahit | B | 3,487,895 | 2,429,981,216 | 452 | 36,251 | 912 |
| *Li. durhamii* | Megahit | C | 3,189,279 | 1,984,519,170 | 382 | 216,289 | 747 |
| *Cq. hermanoi* | Megahit | Zpp2 | 4,306,268 | 2,319,591,732 | 283 | 44,771 | 692 |
| *Cq. albicosta* | - | - | - | - | - | - | - |

**Table 4.** Number of specimens per pool of mosquitoes.

| Pool name | Mosquito species | Number of specimens pooled |
|---|---|---|
| P10 | *Ma. titillans* | 37 |
| P8 | *Ma. wilsoni* | 26 |
| P7 | *Ae. albopictus* | 25 |
| P4 | *Ps. ferox* | 22 |
| P3 | *Ae. scapularis* | 18 |
| P6 | *Cq. chrysonotum* | 15 |
| P2 | *Cq. venezuelensis* | 15 |
| P5 | *Li. durhamii* | 15 |
| P9 | *Cq. hermanoi* | 13 |
| P1 | *Cq. albicosta* | 8 |

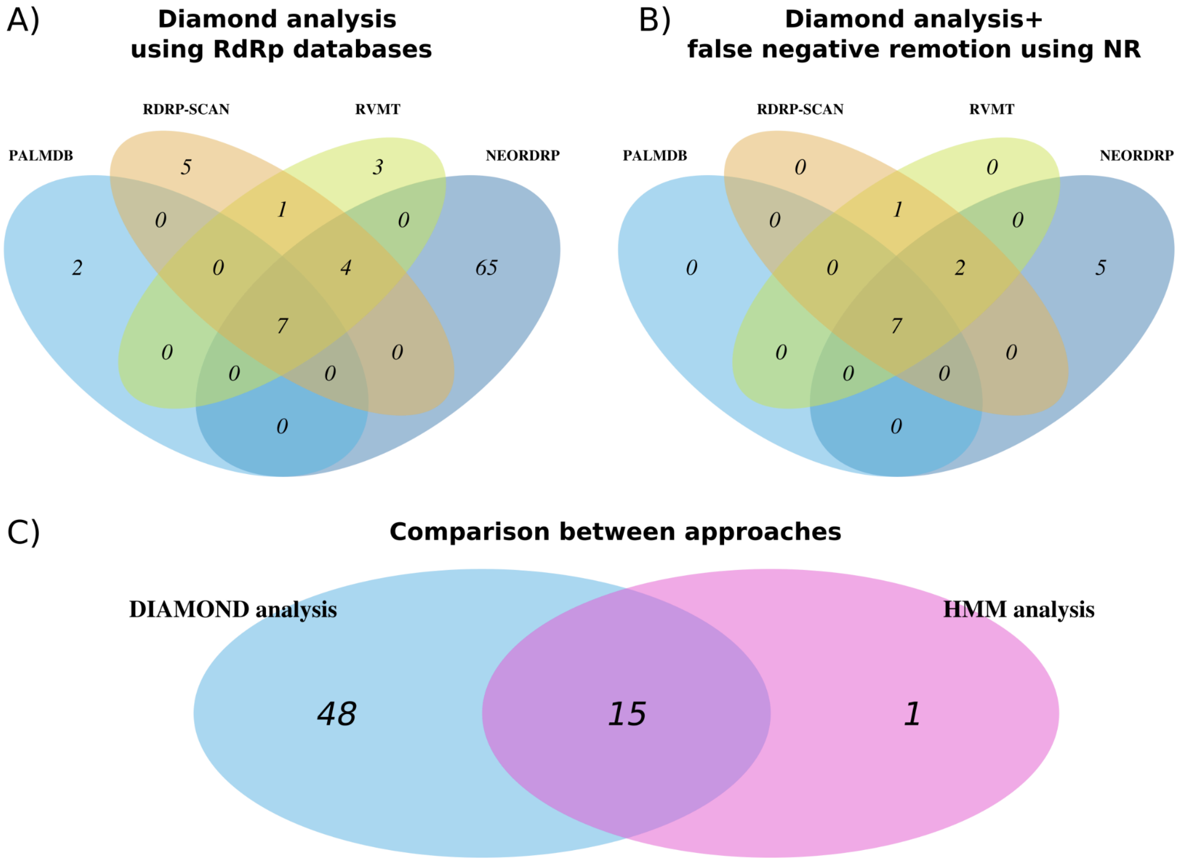


**Figure 4.** General comparison across approaches for viral identification. A) Venn diagram showing the overlap of viral contigs identified by each database used on DIAMOND analysis. B) Venn diagram showing the overlap of viral contigs identified by each database used on DIAMOND analysis after false positive hits remotion.

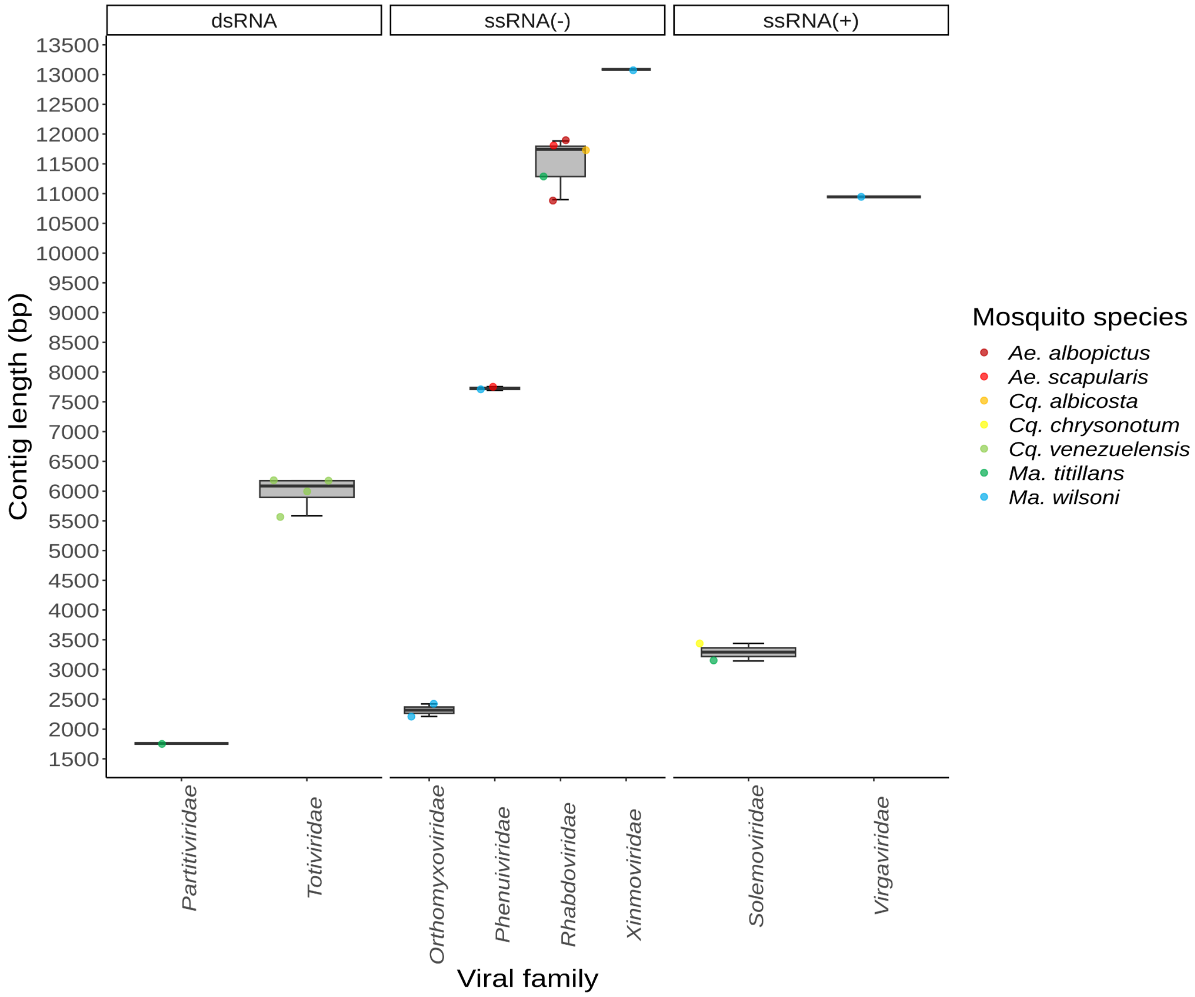


**Figure 5.** Contig length distribution of full-length viral sequences with RdRp markers across mosquito species. The boxplots show the distribution of contig lengths for each viral family identified in mosquitoes. Colored dots represent individual contigs identified in the viromes. Viral families with full-length sequences identified are shown in bold.

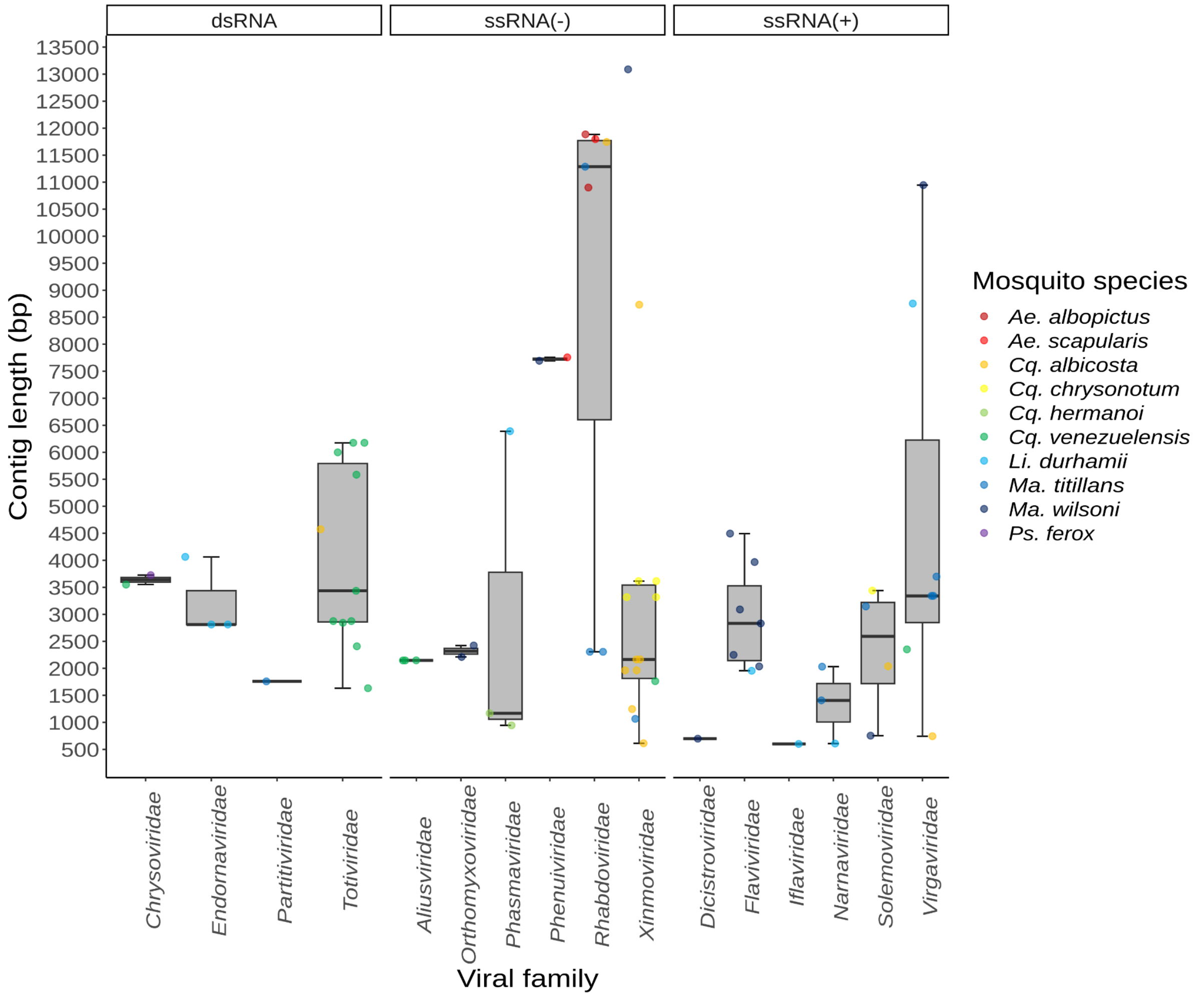


**Figure 6.** Contig length distribution of viral sequences with RdRp markers across mosquito species. The boxplots show the distribution of contig lengths for each viral family identified in mosquitoes. Colored dots represent individual contigs identified in the viromes.

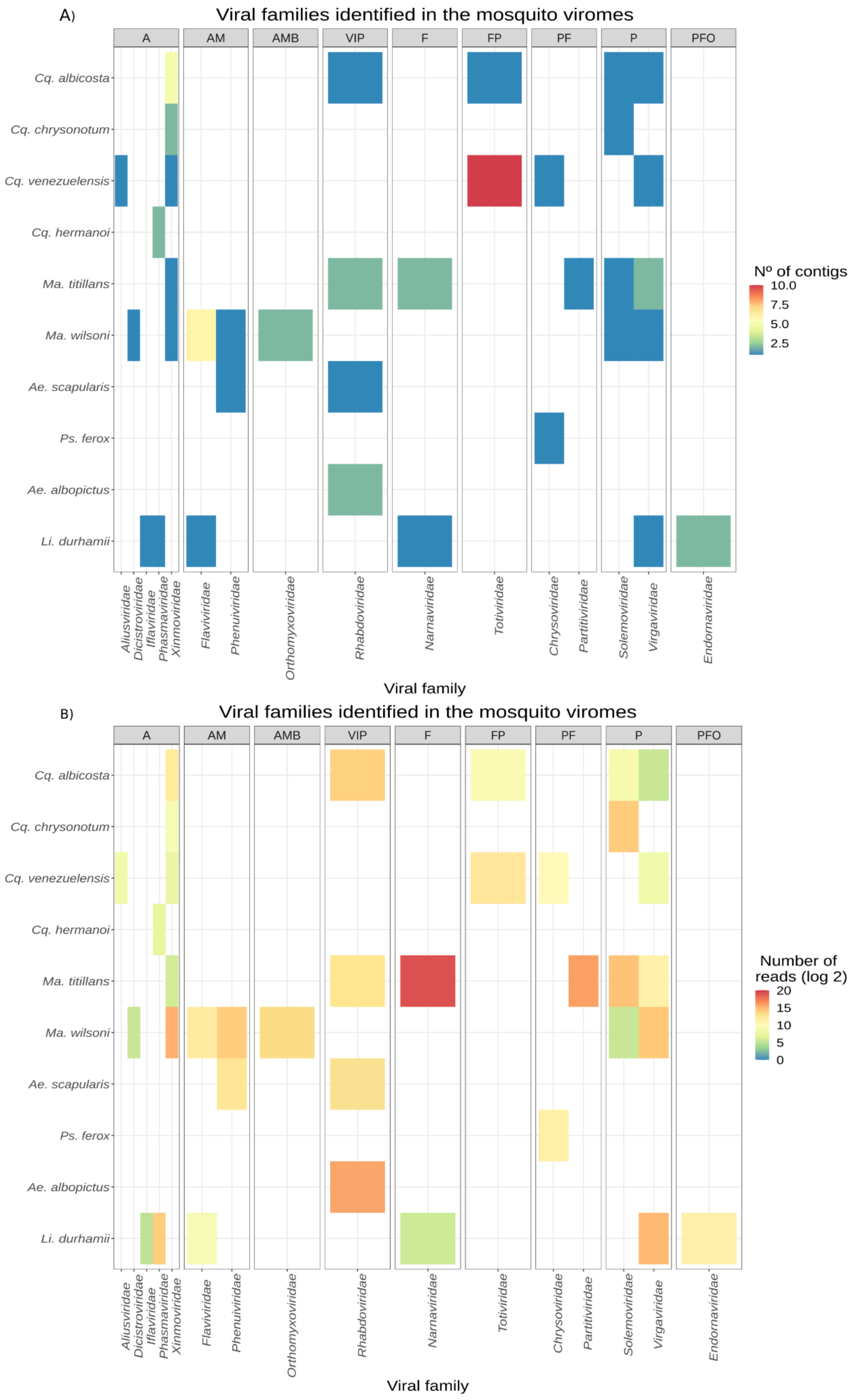


**Figure 7**.Heat maps of viruses identified across mosquito viromes. The grids are different groups of hosts infected by each viral family according to ICTV (www.talk.ictvonline.org/) and ViralZone ([www.viralzone.expasy.org/](www.viralzone.expasy.org/)). A) Heatmap showing the number of viral

contigs identified by mosquito species and viral family. B) Heatmap showing the number of reads identified by mosquito species and viral family. A - Arthropoda, IF - Insects or Fishes, AM - Arthropoda and mammals, AMB - Arthropods, mammals and birds, MBF - Mammals, birds and fishes, VIP - Vertebrates, invertebrates and plants, F - Fungi, FP - Fungi and protozoa, P - Plants, PF - Plants and Fungi, PFO - Plants, fungi and oomycetes.

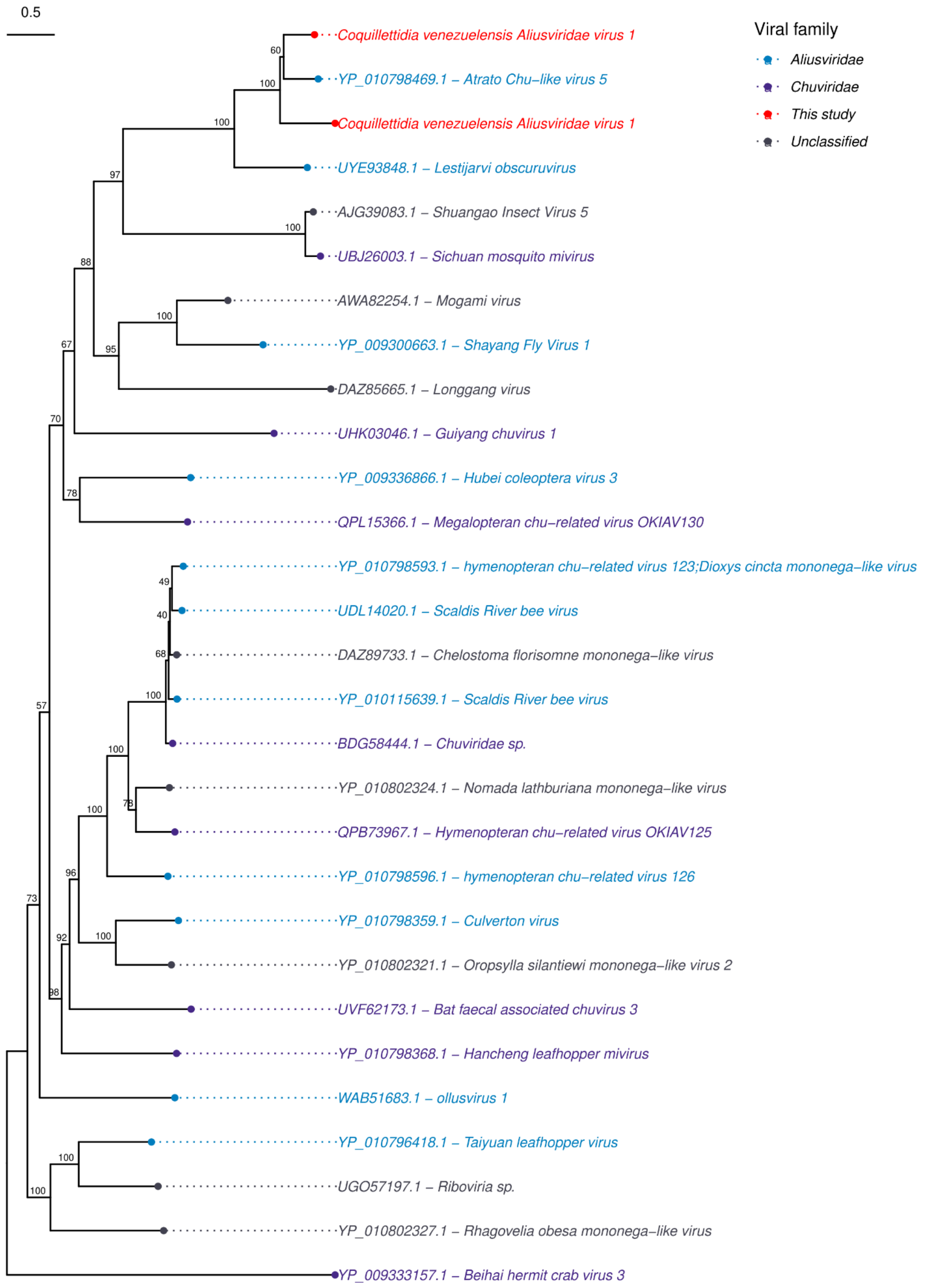


**Figure 8.** Phylogenetic tree of Jungchuvirales order (*Aliusviridae* and *Chuviridae*). The phylogenetic tree included twenty nine sequences and was reconstructed based on a 2,454 aligned amino acid sites while our sequences showed between 118 and 125 aligned amino

acid sites after trimming representing the RdRp sequences and analyzed on IQ-TREE2.0 performing the ultrafast bootstrapping with 1,000 replicates using the LG+F+I+G4 as evolutionary model. The tree was set as the midpoint root. The colors represent the different viral families of viruses used in the analysis.

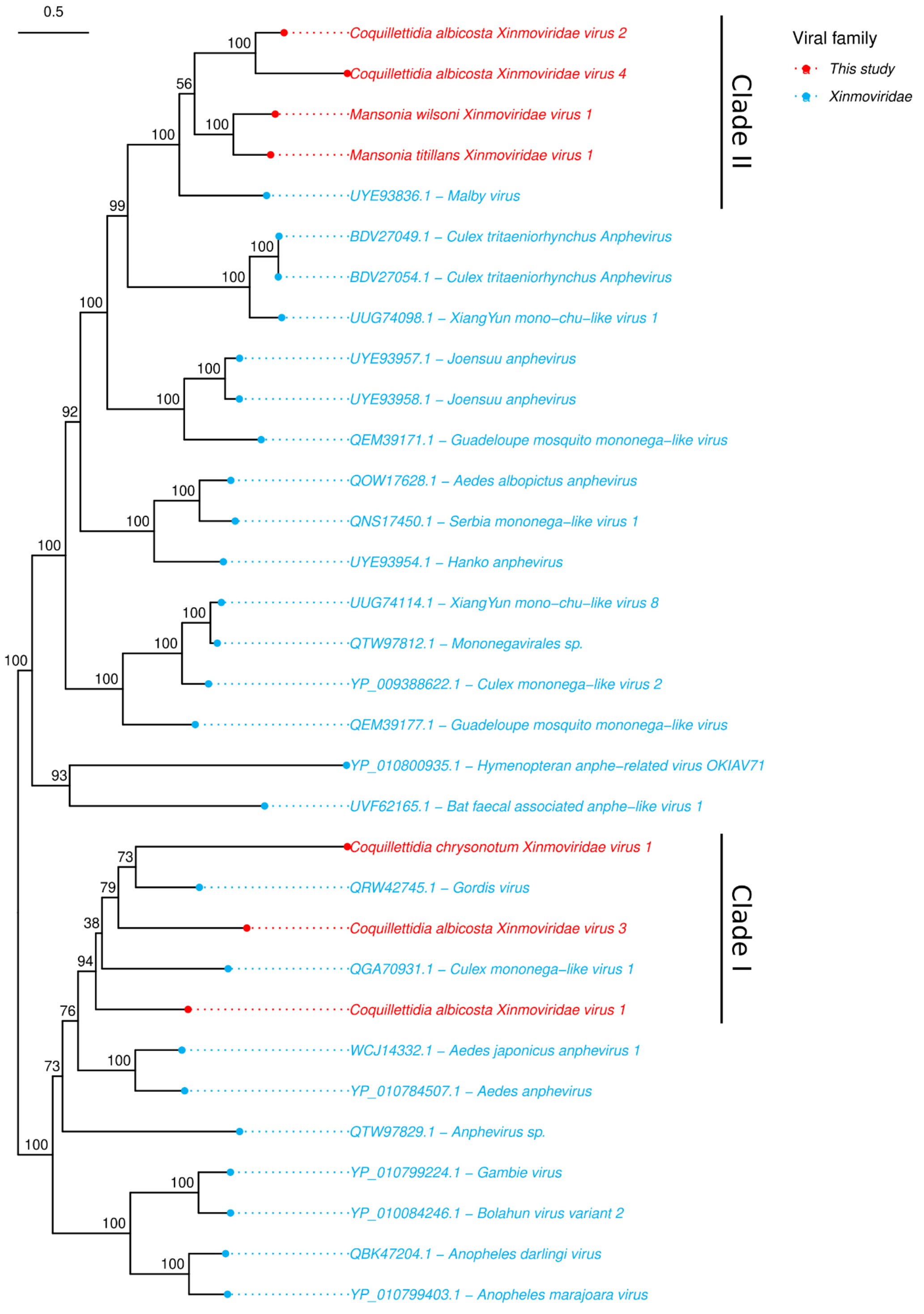


**Figure 9.** Phylogenetic tree of the *Xinmoviridae* family. The phylogenetic tree included thirty two sequences and was reconstructed based on a 1,998 aligned amino acid sites while our sequences ranged from 125 to 1,993 aligned amino acid sites after trimming representing the

RdRp sequences and analyzed on IQ-TREE2.0 performing the ultrafast bootstrapping with 1,000 replicates using the the  LG+F+I+G4 as evolutionary model. The tree was set as the midpoint root. The colors represent the different viral families of viruses used in the analysis.

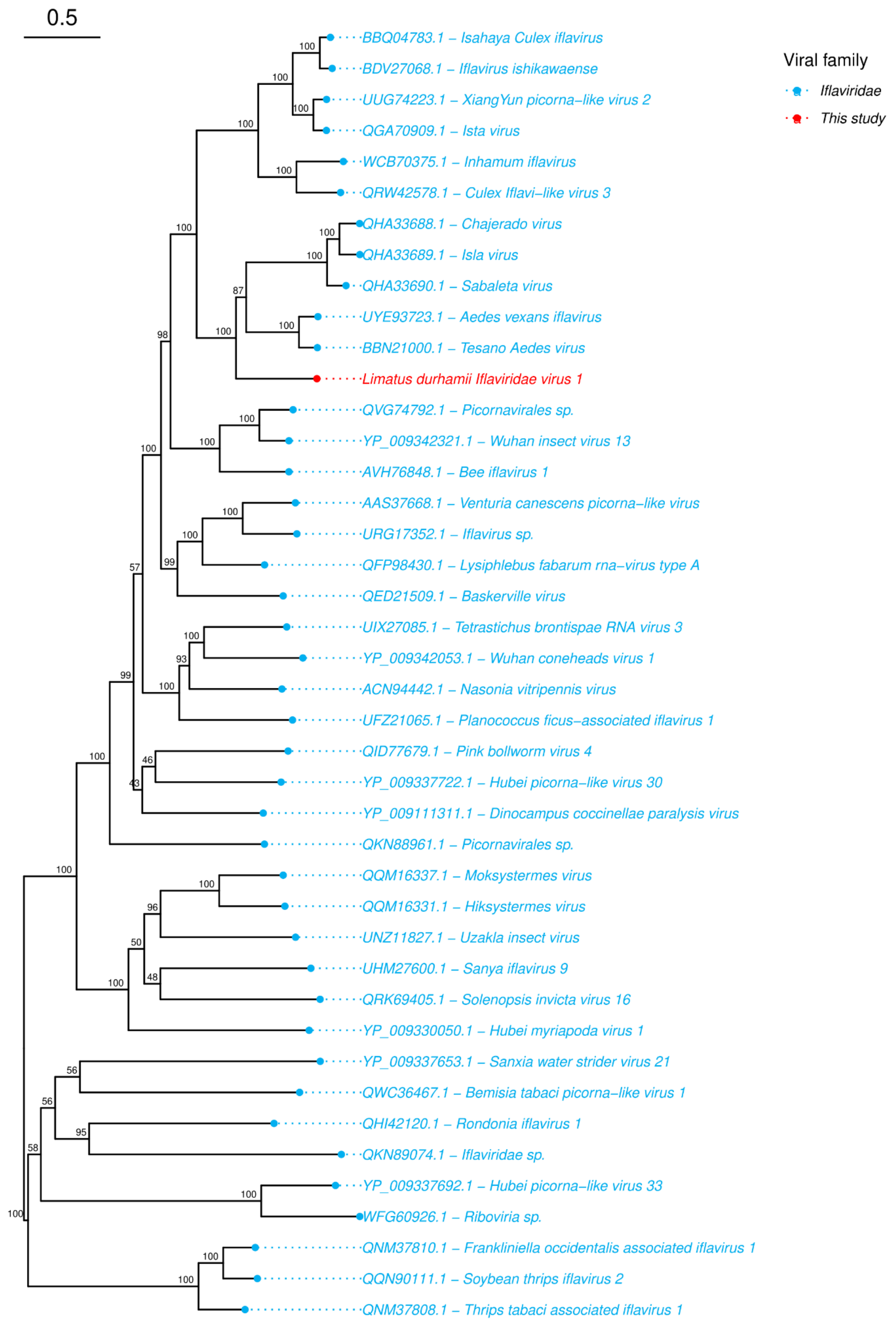


**Figure 10.** Phylogenetic tree of *Iflaviridae* family. The phylogenetic tree included forty five sequences and was reconstructed based on a 2,664 aligned amino acid incuding our sequence of 199 aligned amino acid sites representing the RdRp sequence and analyzed on

IQ-TREE2.0 performing the ultrafast bootstrapping with 1,000 replicates using the the Q.pfam+F+I+G4 as evolutionary model. The tree was set as the midpoint root. The colors represent the different viral families of viruses used in the analysis.

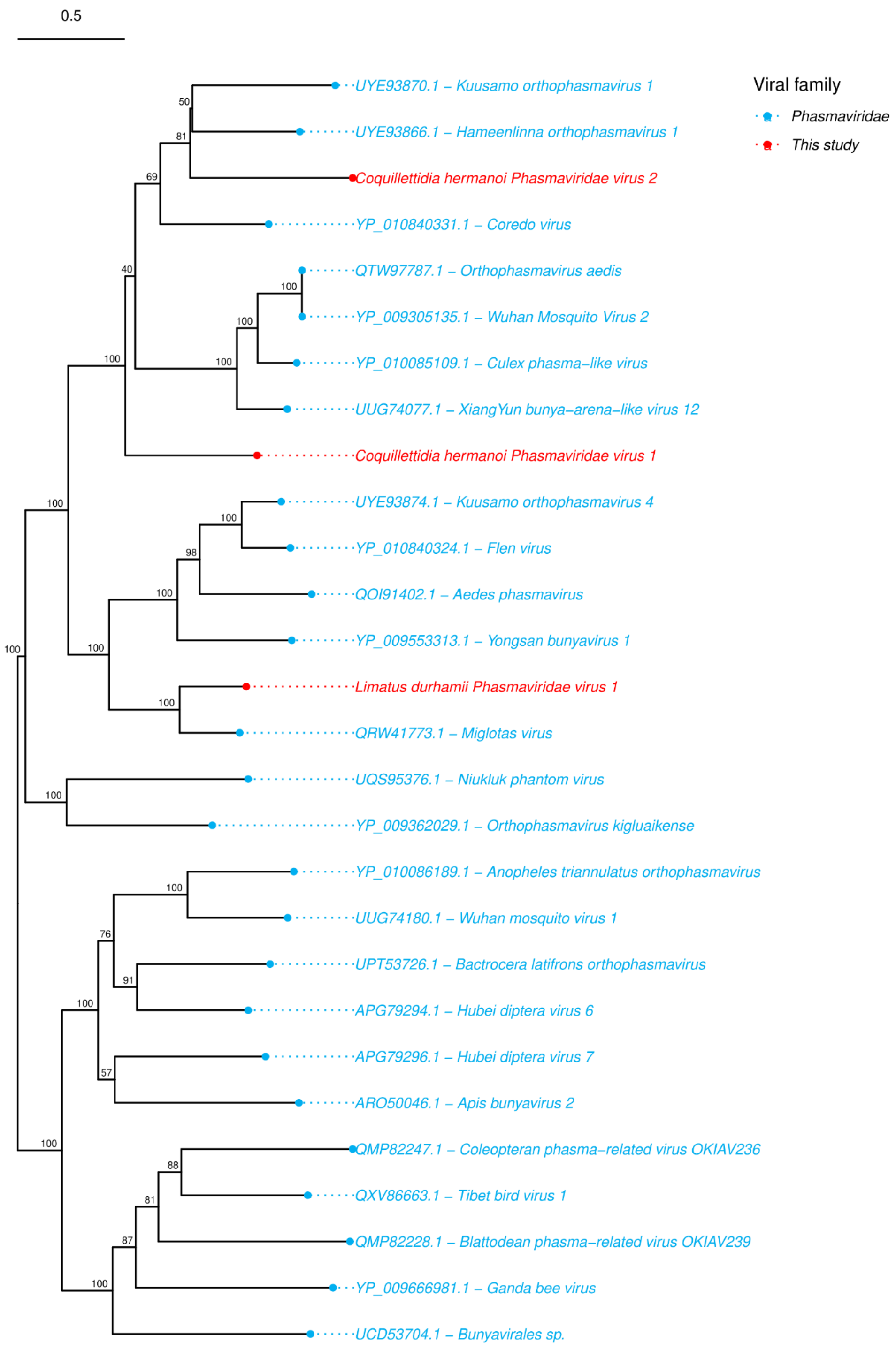


**Figure 11.** Phylogenetic tree of *Phasmaviridae* family. The phylogenetic tree included twenty eight sequences and was reconstructed based on a 2,083 aligned amino acid sites while our sequences showed between 312 and 2,074 aligned amino acid sites after trimming representing the RdRp sequences and analyzed on IQ-TREE2.0 performing the ultrafast

bootstrapping with 1,000 replicates using the the Q.pfam+F+I+G4 as evolutionary model. The tree was set as the midpoint root. The colors represent the different viral families of viruses used in the analysis.

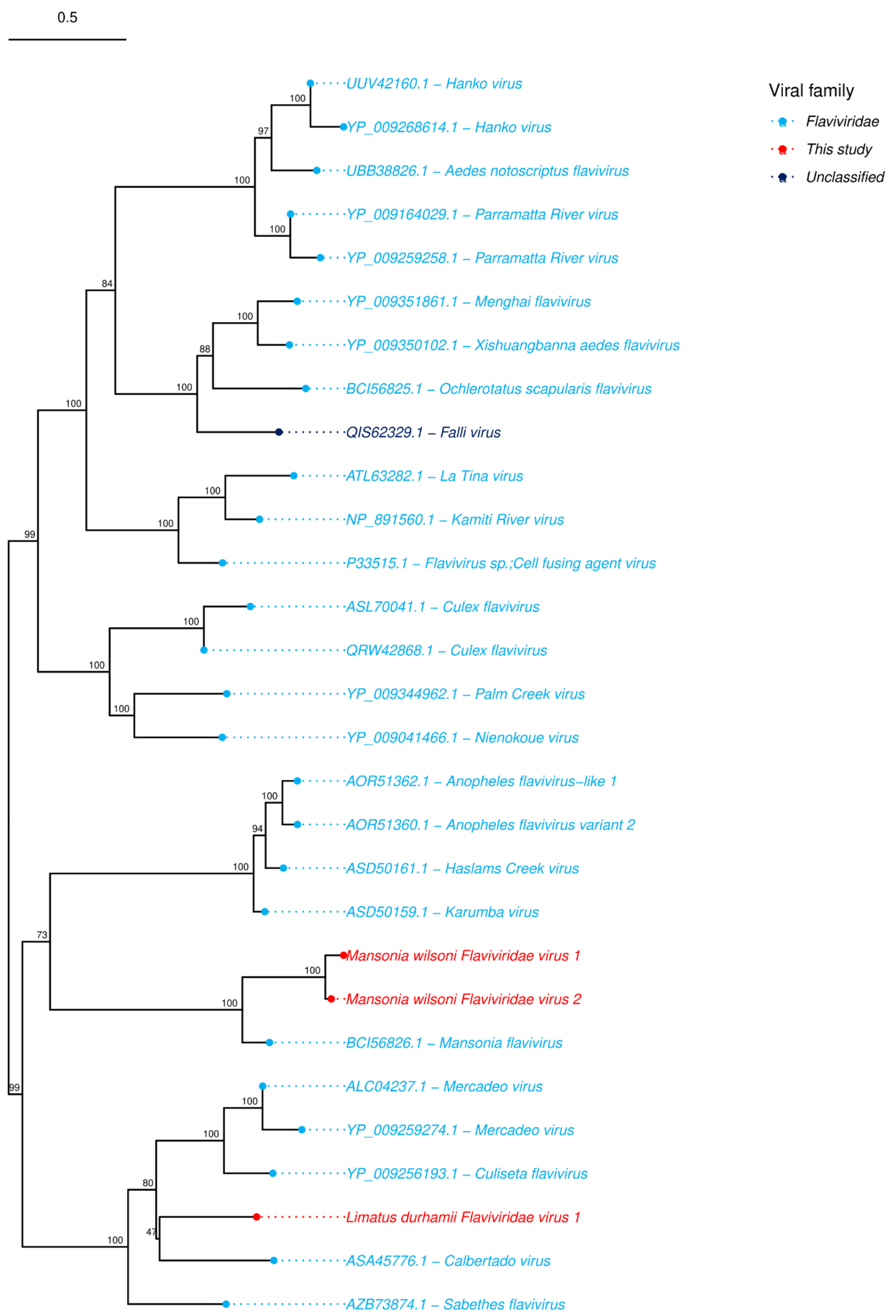


**Figure 12.** Phylogenetic tree of the *Flaviviridae* family. The phylogenetic tree included twenty nine sequences and was reconstructed based on a 3,302 aligned amino acid sites while our sequences ranged from 126 to 212 aligned amino acid sites representing fragments of polyprotein sequences and analyzed on IQ-TREE2.0 performing the ultrafast bootstrapping

with 1,000 replicates using the the LG+F+I+G4 as evolutionary model. The tree was set as the midpoint root. The colors represent the different viral families of viruses used in the analysis.

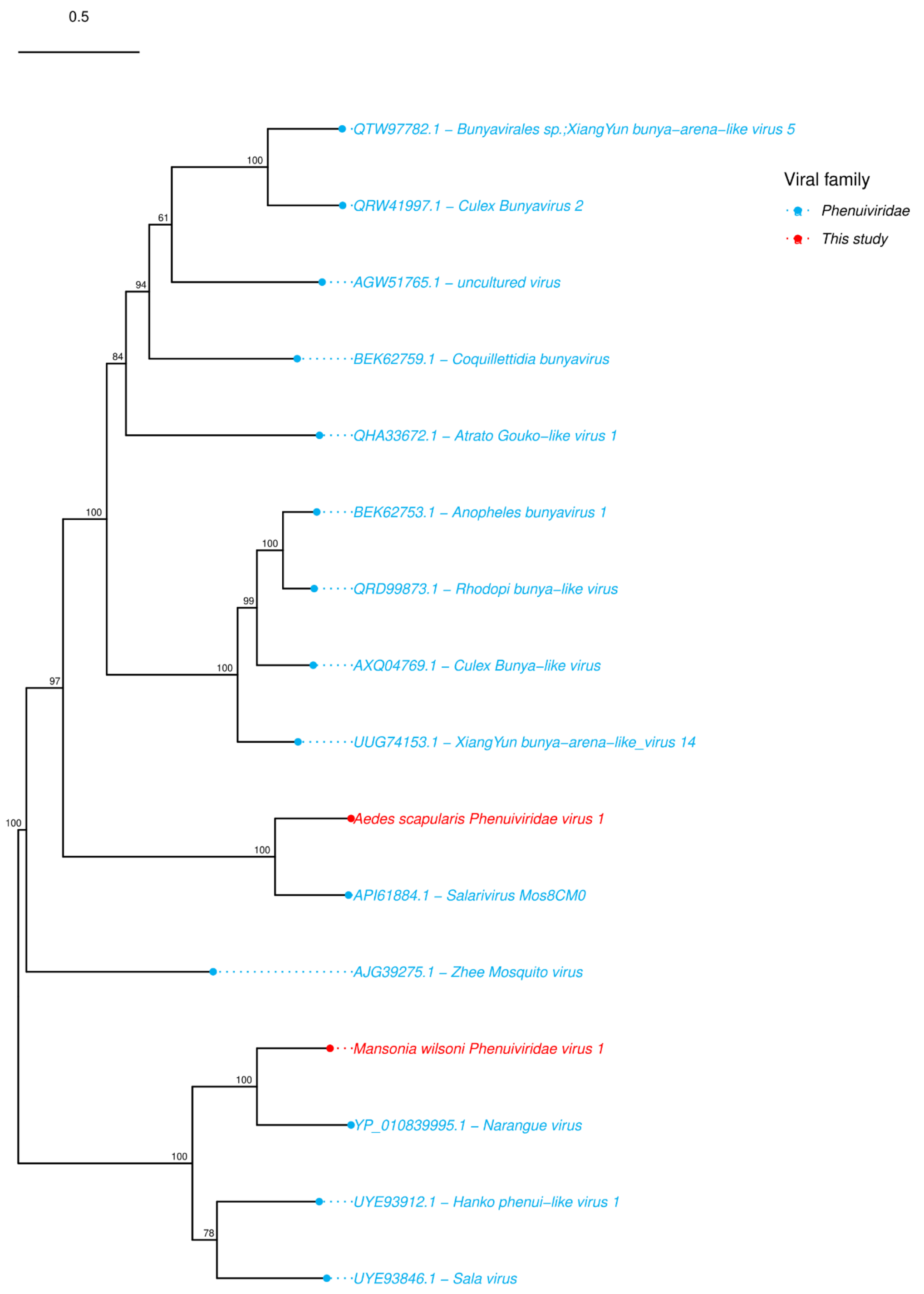


**Figure 13.** Phylogenetic tree of *Phenuiviridae* family. The phylogenetic tree included twenty eight sequences and was reconstructed based on a 2,450 aligned amino acid sites while our sequences ranged from 2411 to 2426 aligned amino acid sites representing the RdRp sequences and analyzed on IQ-TREE2.0 performing the ultrafast bootstrapping with 1,000

replicates using the the LG+F+I+G4 as evolutionary model. The tree was set as the midpoint root. The colors represent the different viral families of viruses used in the analysis.

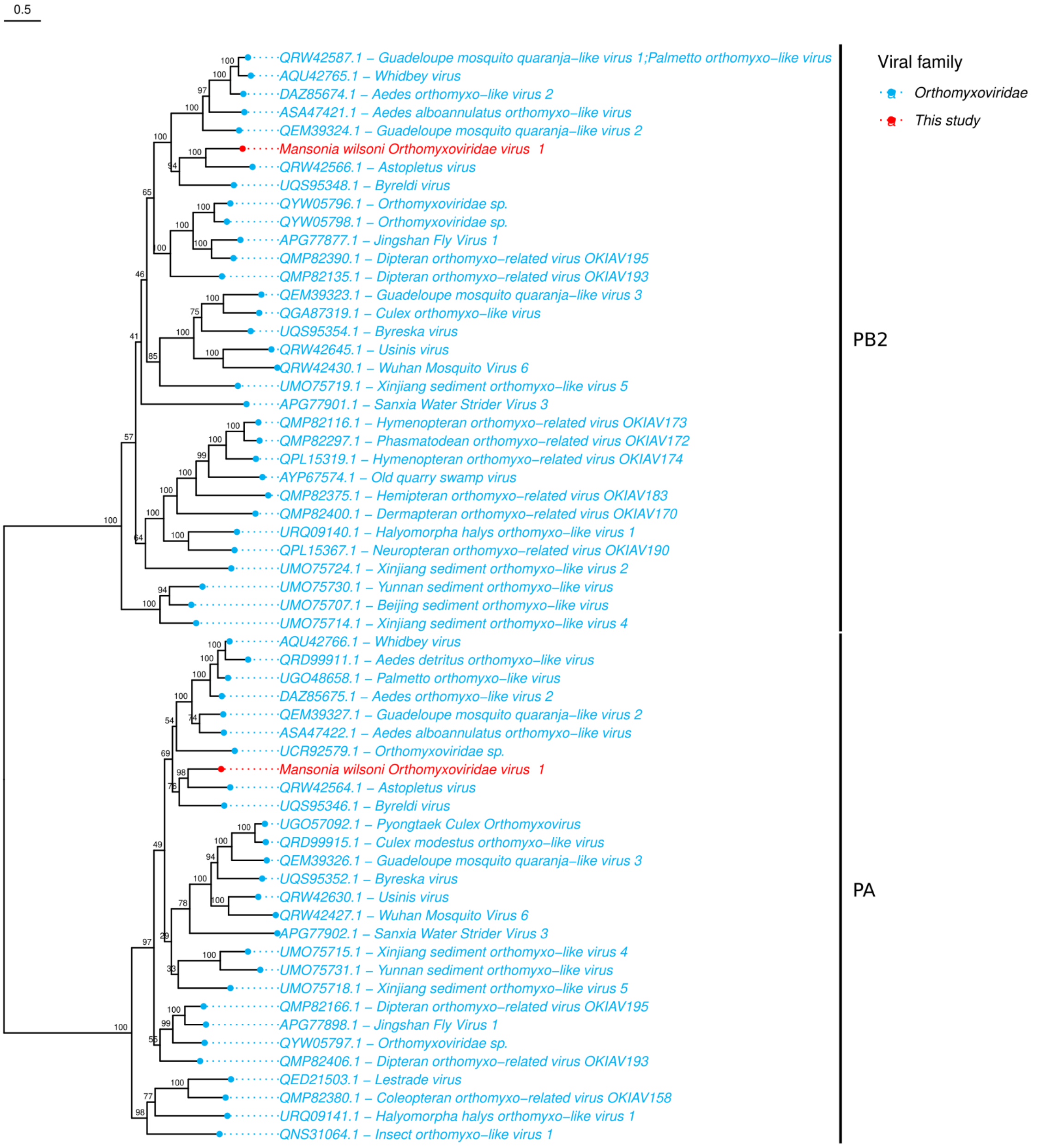


**Figure 14.** Phylogenetic tree of the *Orthomyxoviridae* family. The phylogenetic tree included ninety sequences and was reconstructed based on a 708 aligned amino acid sites while our sequences ranged from 560 to 691 aligned amino acid sites representing the RdRp sequences (PA and PB2) and analyzed on IQ-TREE2.0 performing the ultrafast bootstrapping with 1,000 replicates using the the rtREV+F+G4 as evolutionary model. The tree was set as the midpoint root. The colors represent the different viral families of viruses used in the analysis.

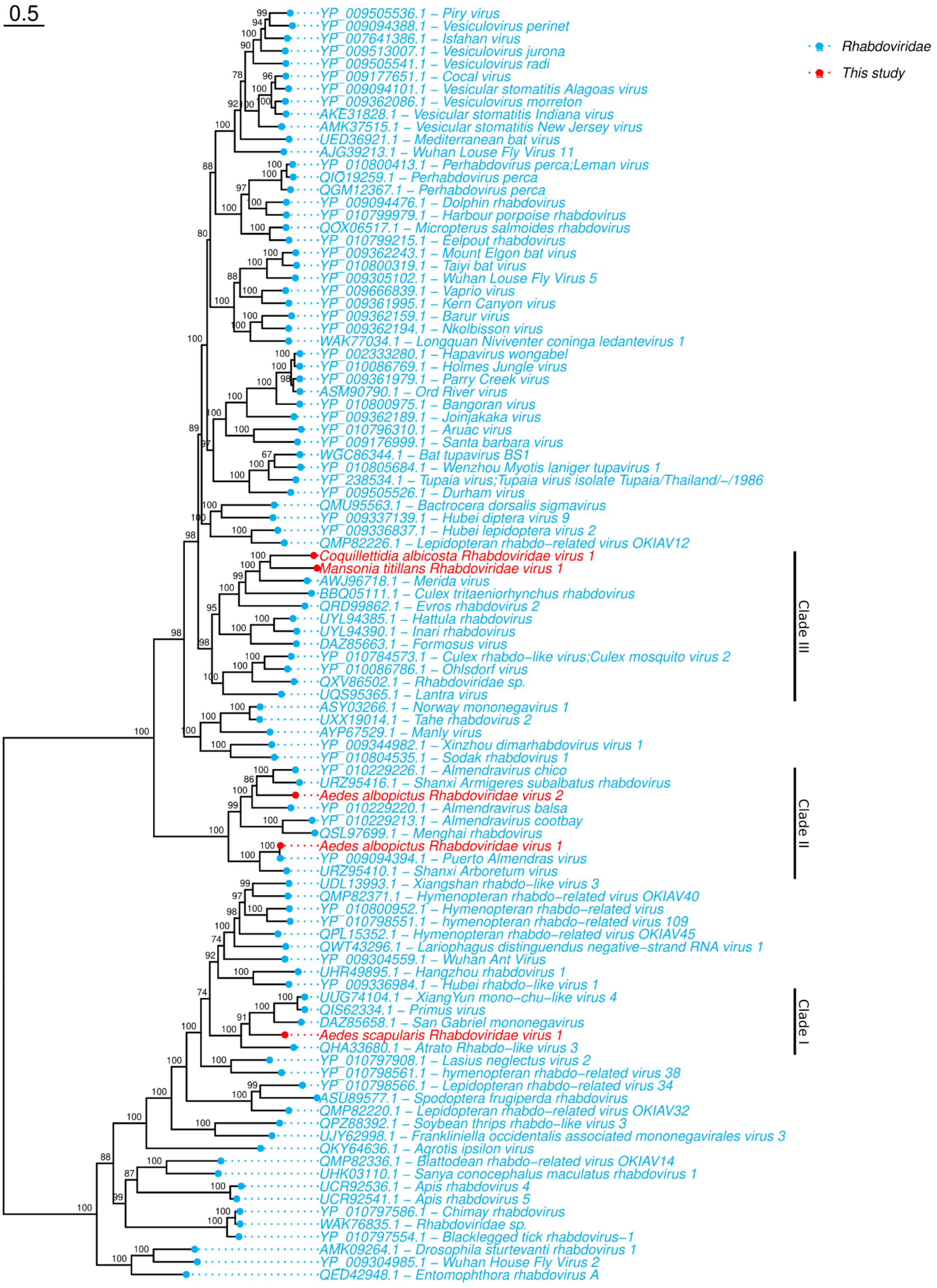


**Figure 15.** Phylogenetic tree of the *Rhabdoviridae* family. The phylogenetic tree included one hundred one sequences and was reconstructed based on a 2,101 aligned amino acid sites while our sequences ranged from 1,851 to 2,071 aligned amino acid sites representing the RdRp sequences and analyzed on IQ-TREE2.0 performing the ultrafast bootstrapping with 1,000 replicates using the the Q.pfam+F+I+G4 as evolutionary model. The tree was set as the midpoint root. The colors represent the different viral families of viruses used in the analysis.

### Supplementary results

#### Fungi, plant, protozoa and oomycetes viruses

For viruses with host assignment in fungi, plant, protozoa and oomycetes we reconstructed a phylogenetic trees for *Narnaviridae* (**Figure 16**), *Totiviridae* (**Figure 17**), *Chrysoviridae* (**Figure 18**), *Partitiviridae* (**Figure 19**), *Solemoviridae* (**Figure 20**) and *Virgaviridae* (**Figure 21**).

For Narnaviridae tree we analyzed 34 sequences based on an alignment of 798 amino acid sites (**Figure 16**). Only one sequence identified here was included representing a *Narnaviridae* sequence from *Ma. titillans* that showed 546 aligned amino acid sites. This sequence was placed into a clade together with Hubei mosquito virus 3 (UUG74035.1) and Hubei narna−like virus 14 (APG77221.1). However, our sequence was more closely of the first one virus that was identified in mosquitoes from China. For *Totiviridae* we analyzed 22 sequences based on 874 aligned amino acid sites and we included three sequences, two *Cq. venezuelensis* and one from *Cq. albicosta* that showed 618 to 863 aligned amino acid sites and clustered all together (**Figure 17**). For *Chrysoviridae* tree we analyzed 29 sequences based on an alignment of 1,102 sites. Two sequences were identified here, representing complete segments with 1,075 and 1,091 aligned amino acid sites for *Cq. venezuelensis* and *Ps. ferox* respectively (**Figure 18**). These sequences were placed into a high supported clade (UFboot of 99) together with Hanko alphachrysovirus (UUV42195.1), Keturi virus (QRW42852.1), Enontekio alphachrysovirus (UUV42191.1) and Lestijarvi alphachrysovirus (UUV42326.1 and UUV42200.1). *Psorophora ferox* Chrysoviridae virus 1 was placed as basal branch in relation to the remaining sequences. While, *Coquillettidia venezuelensis* Chrysoviridae virus 1 was closely with Keturi virus (QRW42852.1). For *Partitiviridae* we analyzed 28 sequences based on 533 aligned amino acid sites and we included one sequence from *Ma. titillans* that showed 521 aligned amino acid sites (**Figure 19**). This sequence was placed as a basal branch from a clade containing Hattula partiti−like virus (UUV42351.1) and Hubei partiti−like virus 22 (BDV27034.1).

For Solemoviridae we analyzed 38 sequences based on 422 aligned amino acid sites and we included 4 sequences identified here representing *Ma. titillans* (N=1), *Cq. albicosta* (N=1) and *Cq. chrysonotum* (N=1) and *Ma. wilsoni* (N=1 )(**Figure 20**). Our sequences were placed into three distinct clades. The first one clustered one sequence from *Ma. wilsoni* (*Mansonia wilsoni Solemoviridae* virus 1) into basal clade in relation to all phylogeny. The viral sequence from *Cq. chrysonotum* (Coquillettidia chrysonotum Solemoviridae virus 1) clustered with Atrato Sobemo−like virus 6 (QHA33729.1). The viral sequences for *Ma. titillans and Cq. chrysonotum* clustererd together with Atrato Sobemo−like virus 5 (QHA33869.1) identified in *Cq. venezuelensis* from Colombia.

For the Virgaviridae tree we analized 61 sequences based on 2,277 aligned amino acid sites and 5 sequences of them were identified here representing *Ma. titillans* (N=1), *Cq. venezuelensis* (N=1), *Cq. albicosta* (N=1), *Li. durhamii* (N=1) and *Ma. wilsoni* (N=1). Our sequences ranged from 126 up to 2,043 aligned amino acid sites (**Figure 21**). In general, our sequences were placed into 3 clades. The sequences from *Cq. venezuelensis, Cq. albicosta* and *Li. durhamii* clustered together Atrato Virga−like virus 2 (QHA33738.1). However, this virus was closely with *Coquillettidia venezuelensis Virgaviridae* virus 1. *Mansonia wilsoni Virgaviridae virus 1* sequence clustered together Atrato Virga−like virus 5 (QHA33754.1) and the high amino acid identity of RdRp (95%) suggest that are the same virus. Another clade identified was composed by the sequences from *Ma. titillans* that clustered together with Atrato Virga−like virus 7 (QHA33782.1) identified in *Ps. albipes* from Colombia.

**Unclassified sequences**

One sequence from those 64 identified previously was analyzed phylogeneticly and we reconstructed a tree comprising 35 sequences including sequences from NCBI assigned in *Adintoviridae, Alloherpesviridae, Allomimiviridae, Baculoviridae, Mimiviridae, Peribunyaviridae, Phasmaviridae* and *Phycodnaviridae* (**Figure 21**). The phylogenetic tree was reconstructed based on 742 aligned amino acid sites and our sequence showed 162 aligned amino acid sites representing an unclassified sequence for *Cq. hermanoi* (*Coquillettidia hermanoi Unclassified* virus 1). This particular sequence was placed within a clade consisting of sequences from the *Phasmaviridae* and other unclassified sequences. However, our analysis did not show a high UFboot value to support an accurate taxonomic classification of this sequence that remains as assigned as unclassified.

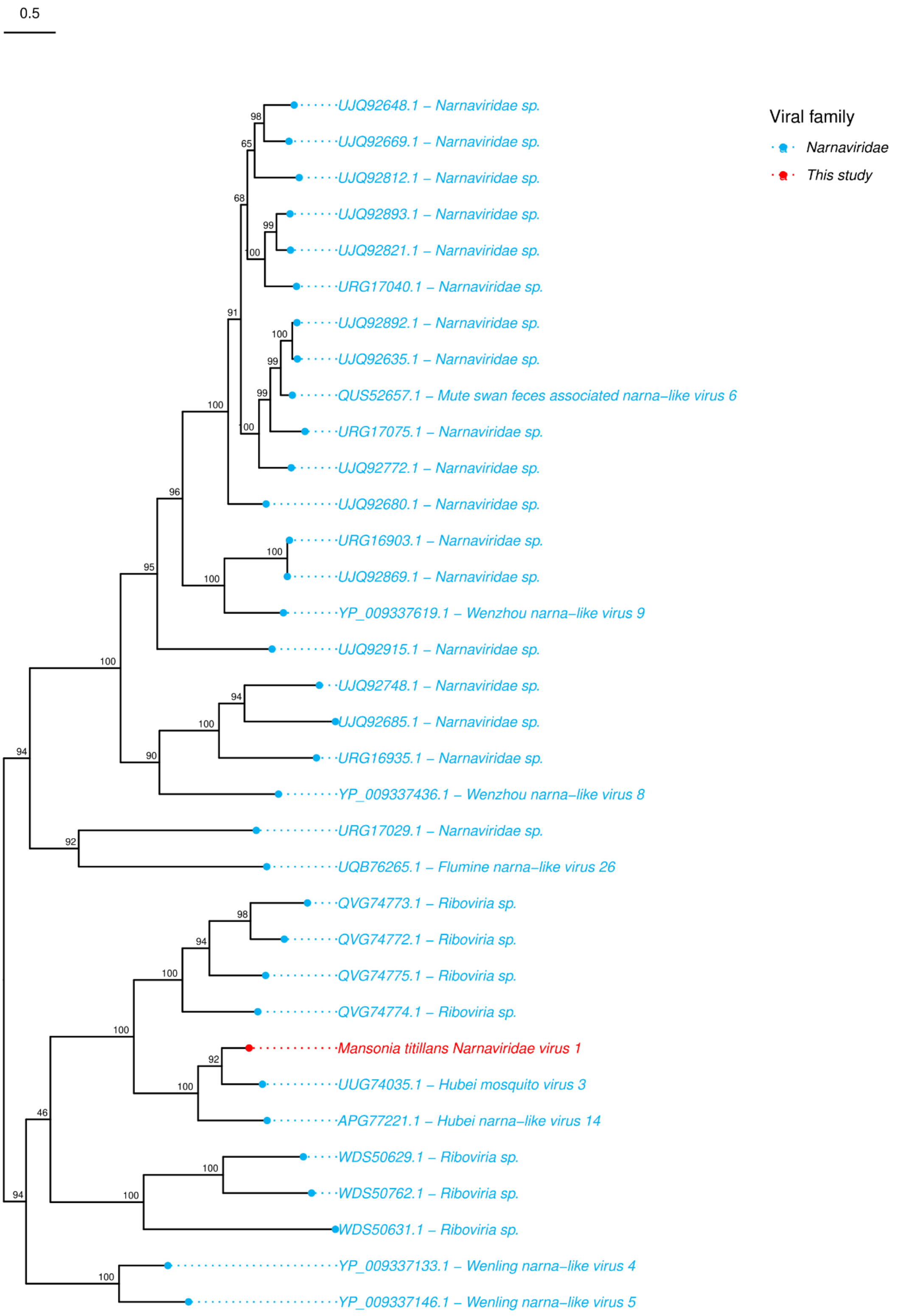


**Figure 16.** Phylogenetic tree of *Narnaviridae* family. The phylogenetic tree included thirty four sequences and was reconstructed based on a 798 aligned amino acid sites while our sequence showed 613 aligned amino acid sites representing the RdRp sequences and analyzed on IQ-TREE2.0 performing the ultrafast bootstrapping with 1,000 replicates using the the Q.pfam+F+I+G4 as evolutionary model. The tree was set as the midpoint root. The colors represent the different viral families of viruses used in the analysis.

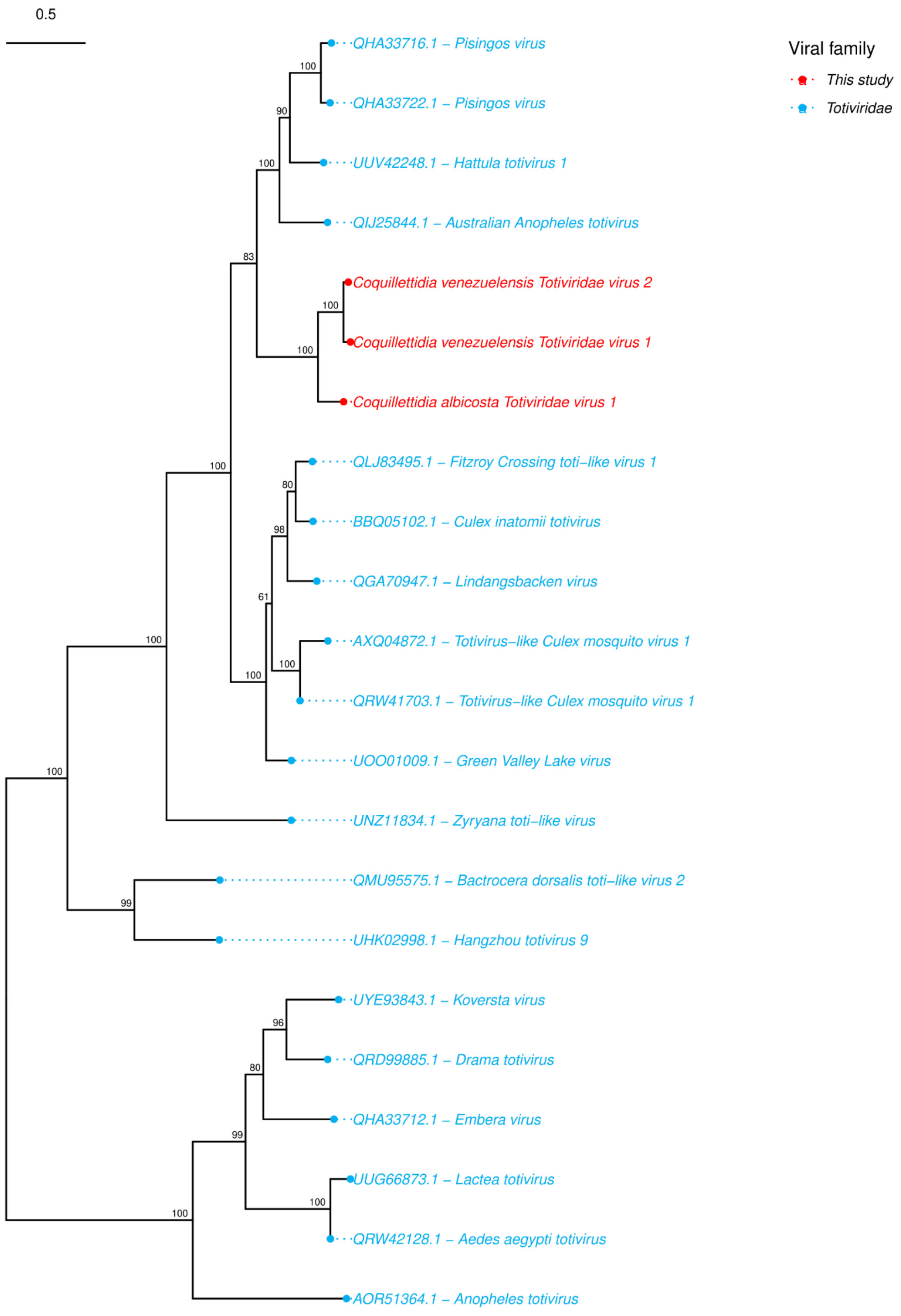


**Figure 17.** Phylogenetic tree of the *Totiviridae* family. The phylogenetic tree included twenty two sequences and was reconstructed based on a 874 aligned amino acid sites while our sequences ranged from 618 to 863 aligned amino acid sites representing the RdRp sequences and analyzed on IQ-TREE2.0 performing the ultrafast bootstrapping with 1,000 replicates using the the LG+F+I+G4 as evolutionary model. The tree was set as the midpoint root. The colors represent the different viral families of viruses used in the analysis.

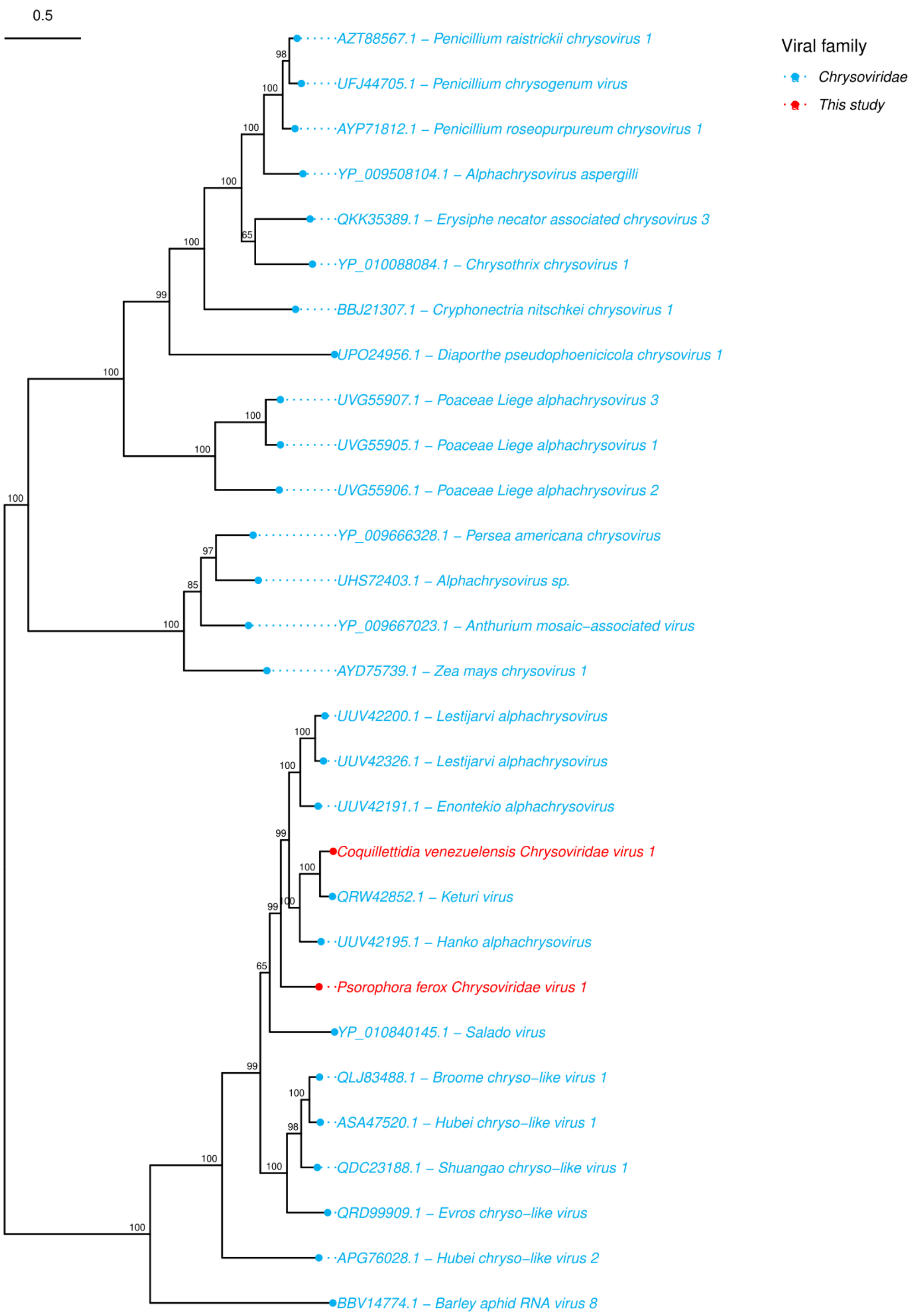


**Figure 18.** Phylogenetic tree of *Chrysoviridae* family. The phylogenetic tree included twenty nine sequences and was reconstructed based on a 1,102 aligned amino acid sites while our sequences ranged from 1,075 to 1,091 aligned amino acid sites representing the RdRp sequences and analyzed on IQ-TREE2.0 performing the ultrafast bootstrapping with 1,000 replicates using the the LG+F+I+G4 as evolutionary model. The tree was set as the midpoint root. The colors represent the different viral families of viruses used in the analysis.

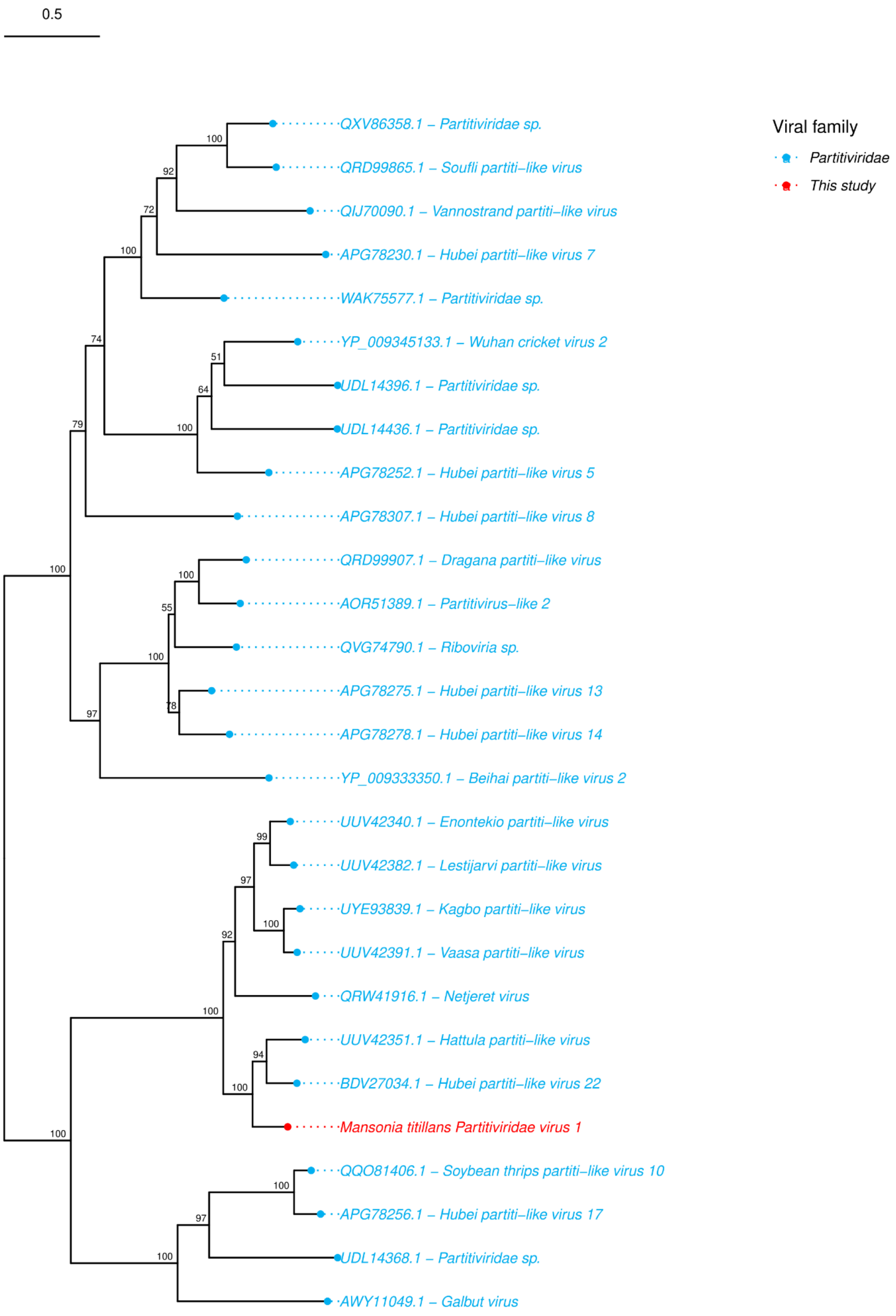


**Figure 19.** Phylogenetic tree of *Partitiviridae* family. The phylogenetic tree included twenty eight sequences and was reconstructed based on a 533 aligned amino acid sites while our sequence showed 521 aligned amino acid sites representing the RdRp sequences and analyzed on IQ-TREE2.0 performing the ultrafast bootstrapping with 1,000 replicates using the the LG+I+G4 as evolutionary model. The tree was set as the midpoint root. The colors represent the different viral families of viruses used in the analysis.

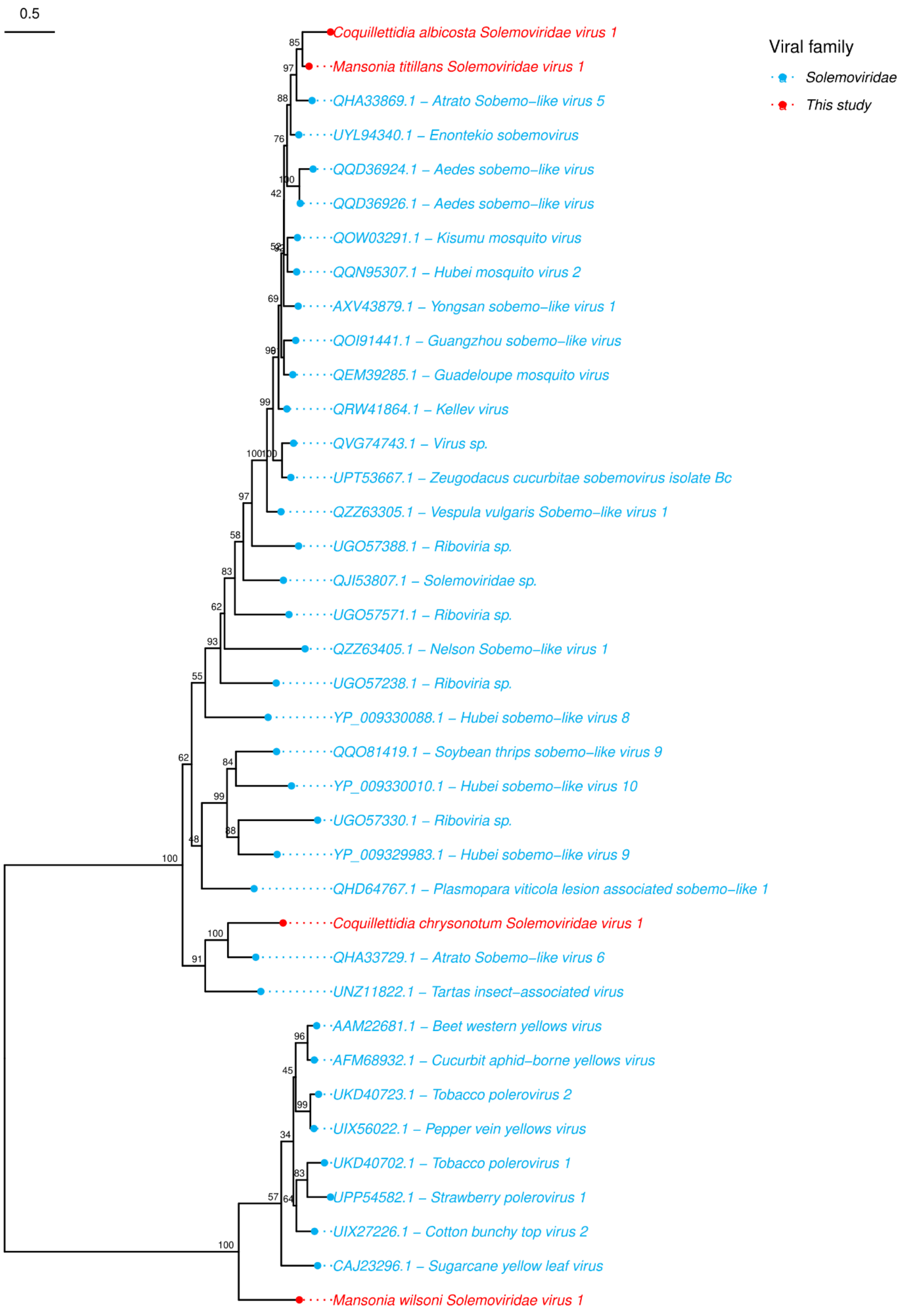


**Figure 20.** Phylogenetic tree of *Solemoviridae* family. The phylogenetic tree included thirty eight sequences and was reconstructed based on a 422 aligned amino acid sites while our sequences ranged from 153 to 351 aligned amino acid sites representing the RdRp sequences and analyzed on IQ-TREE2.0 performing the ultrafast bootstrapping with 1,000 replicates using the the LG+I+G4 as evolutionary model. The tree was set as the midpoint root. The colors represent the different viral families of viruses used in the analysis.

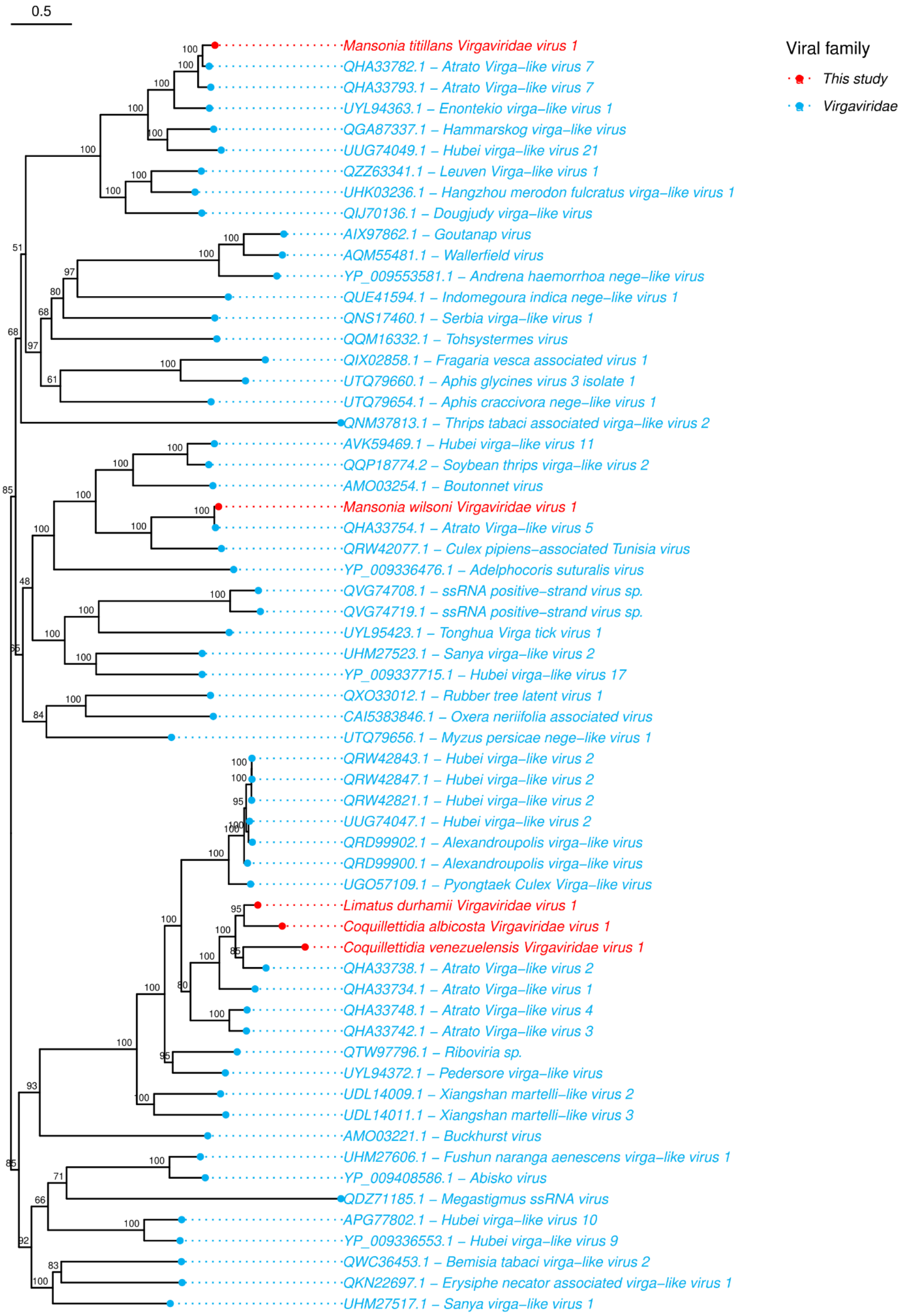


**Figure 21.** Phylogenetic tree of *Vigaviridae* family. The phylogenetic tree included sixty one one sequences and was reconstructed based on a 2,277 aligned amino acid sites while our sequences ranged from 126 to 2,043 aligned amino acid sites representing the RdRp sequences and analyzed on IQ-TREE2.0 performing the ultrafast bootstrapping with 1,000 replicates using the the Q.pfam+F+I+G4 as evolutionary model. The tree was set as the midpoint root. The colors represent the different viral families of viruses used in the analysis.

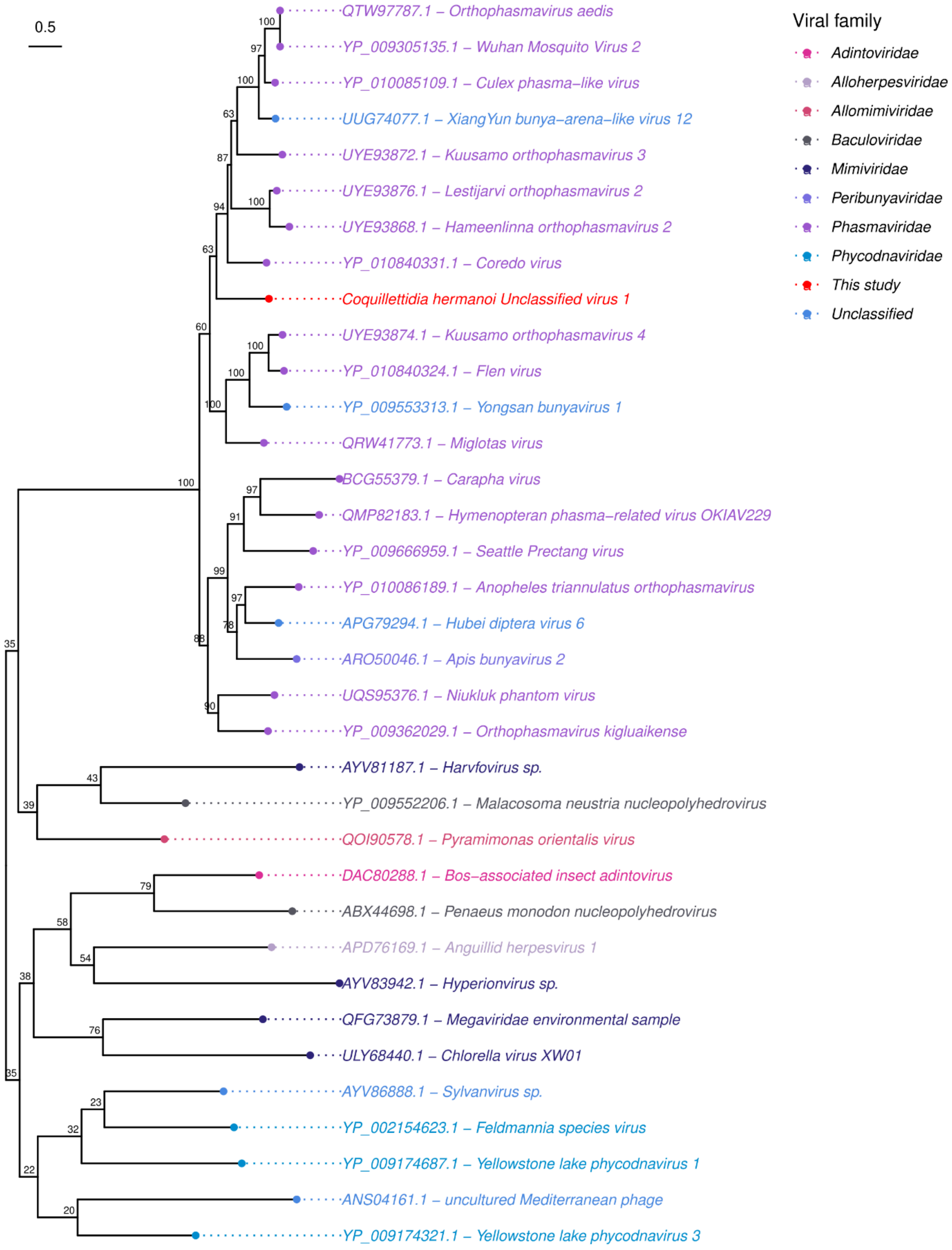


**Figure 22.** Phylogenetic tree of unclassified sequences. The phylogenetic tree included thirty seven sequences and was reconstructed based on a 742 aligned amino acid sites while our sequences ranged from 130 to 232 aligned amino acid sites representing the RdRp sequences

and analyzed on IQ-TREE2.0 performing the ultrafast bootstrapping with 1,000 replicates using the the Q.pfam+F+I+G4 as evolutionary model. The tree was set as the midpoint root. The colors represent the different viral families of viruses used in the analysis.

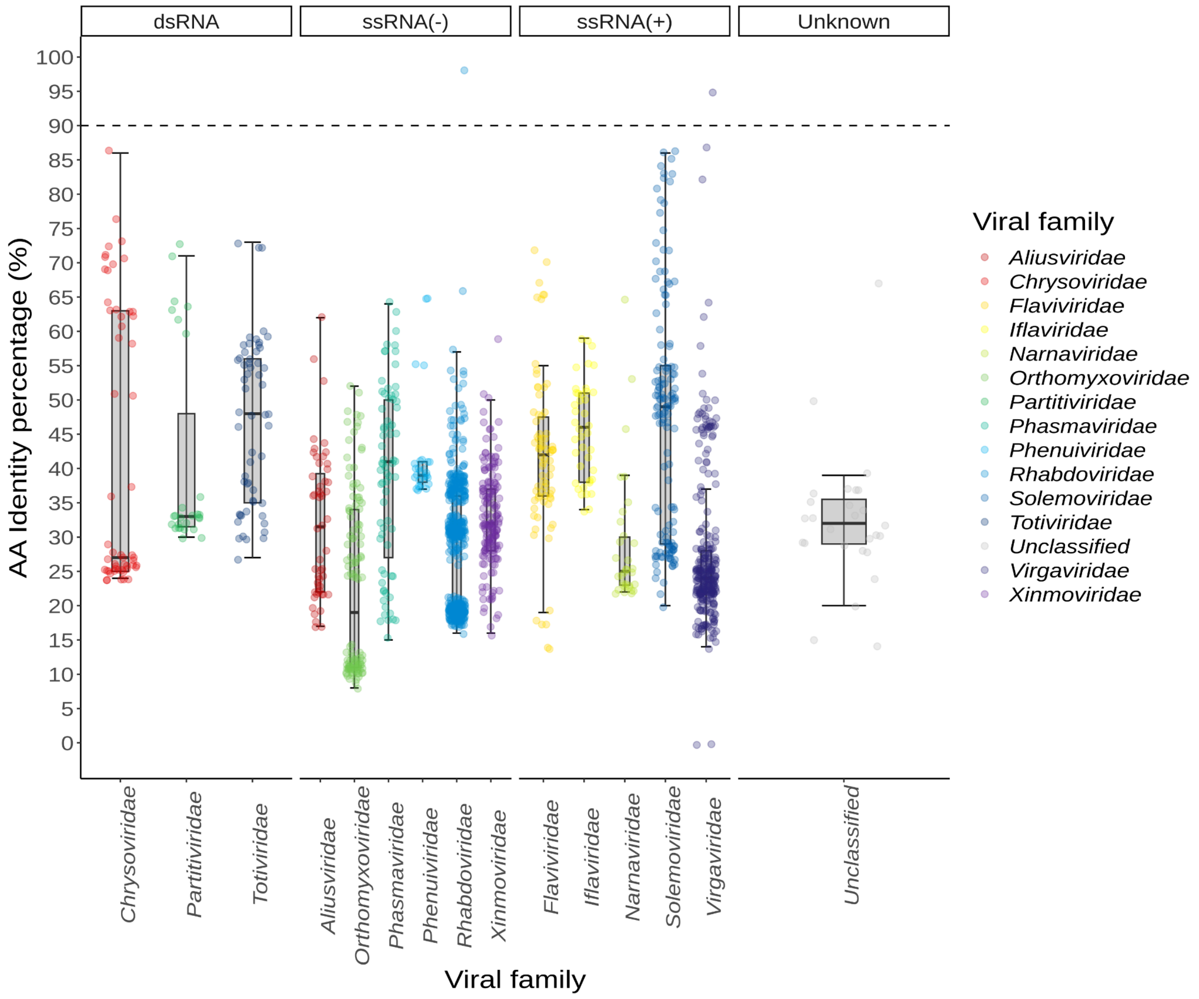


**Figure 23.** Identity percentage of RdRp sequences from identified viral contigs analyzed on Phylogenetic trees against RdRp sequences from NCBI.
